# Distinct Filament Conformation for Receptor-Bound Amyloid-ß from Alzheimer’s Disease Brain

**DOI:** 10.1101/2025.10.10.681740

**Authors:** Mikhail A. Kostylev, Carmen Butan, Graham P. Roseman, Yangyi Liu, Pallavi Gopal, Stephen M. Strittmatter

## Abstract

Alzheimer’s disease is triggered by amyloid-ß, with symptoms linked to synapse loss. Oligomeric amyloid-ß, rather than monomeric or fibrillar amyloid-ß, has been proposed to be the proximate mechanistic cause, but the relevant molecular characteristics have remained unclear. Here, we define a distinct receptor-bound amyloid-ß pool in Alzheimer’s brain by release with a receptor antagonist and purification to homogeneity. Receptor-bound amyloid-ß is ten times more abundant than free unbound amyloid-ß. The amyloid-ß associated with receptor is composed of 65 nm long filaments with prion protein binding at its tips. There is no evidence for an oligomeric Aß state interacting with human brain receptors. Cryo-electron microscopy shows two symmetric S-shaped monomers per filament rung. The tilt between rung monomers, twist along the filament axis, amino terminal conformation and amyloid seeding properties distinguish this structure from plaque-associated amyloid-ß filaments of the same brain. High tip:length ratio is critical for prion protein receptor interaction and synaptic damage. Characterizing receptor-bound amyloid-ß filament provides insight into neuronal dysfunction separate from plaque aggregation.

---

The defining pathology of Alzheimer’s disease (AD) is accumulation of amyloid-ß (Aß) plaques in the brain, with associated neurofibrillary tangles, gliosis and loss of neurons ^1, 2^. Recent anti-Aß antibody studies ^3, 4^, coupled with biomarker progression data ^5^ and rare dominantly inherited causative mutations ^6^, support a causative role for Aß accumulation in triggering the complex disease process ^7^. During the early symptomatic stage, synaptic dysfunction and loss correlate closely with cognitive decline. Soluble, oligomeric forms of Aß (Aßo), as opposed to fibrillary plaque Aß or monomeric Aß, have been postulated to be causative for synaptic phenotypes ^6, 8^. Confusingly, the term Aßo has been applied to many different assemblies of the peptide and the relevant species in the AD brain has remained unclear ^9^. Further, the structure of various Aßo preparations has never been defined, generating uncertainty in the field. In contrast, the structure of fibrillary AD plaque-associated Aß has now been clarified ^10, 11^, and similarities between sporadic and inherited cases have been assessed ^12^.

Reported phenotypes attributed to soluble Aßo include impairment of synaptic plasticity, the triggering of microglial engulfment of synaptic structures, the loss of dendritic spines, the stimulation of phosphorylated Tau accumulation and endolysosomal dysfunction ^6, 8^. A widely studied Aßo preparation of synthetic Aß42 in F12 media, termed amyloid-ß derived diffusible ligands (ADDLs), appear as metastable 500 kDa spherical peptide assemblies ^13^. Globulomers of Aß42 are prepared in SDS and then isolated as stable 12-mers ^14^. From AD brain tissue, freely soluble Aßo species of about 1,000 kDa have been collected after tissue homogenization or soaking of diced brain in buffered saline ^15^. A recent study suggests that at least under certain conditions these preparations contain short Aß filaments that can be sedimented with high centrifugal force ^16^ and that are indistinguishable from insoluble plaque filaments. Thus, the existence of a disease-relevant oligomeric state of Aß is uncertain.

The response of AD brain to the presence of misfolded Aß depends on interaction with cellular components, including specific receptors. A range of cell surface Aß binding sites have been described, with cellular prion protein (PrP^C^) being unique in its specificity for certain species of Aß and discovery by unbiased screening ^17, 18, 19, 20^. The high affinity interaction of Aß with PrP^C^ has a slow off-rate with a half-life of days. In multiple studies, deletion or blockade of PrP^C^ protects the mouse brain from the deleterious effects of Aß accumulation. Specifically, Aß/PrP^C^ complexes acting with mGluR5 as a co-receptor impair long term potentiation in hippocampal slices, trigger Fyn, Pyk2 and eEF2 kinase activity, increase phosphorylated Tau accumulation, disrupt the neuronal transcriptome, recruit C1q to damaged synapses, mediate synapse loss and lead to impaired spatial and object recognition memory in multiple mouse AD models ^21, 22, 23, 24, 25, 26, 27, 28, 29, 30^. The various soluble Aßo assemblies described above each interact with PrP^C^ ^15, 31^. Importantly, binding in the appropriate ratio leads to a hydrogel phase state that sequesters Aß in a bound state with an altered PrP^C^ conformation ^31^. This hydrogel is present in AD brain, and can be disrupted by a polymeric receptor antagonist, termed poly [4-styrenesulfonic acid-co-maleic acid] (PSCMA) ^23, 31^. Here, molecular understanding of Aß interaction with cellular components is advanced by assessing receptor-bound Aß species, distinct from the soluble-free-unbound pool and the insoluble-plaque pool of Aß. Extraction and purification of receptor-bound Aß allowed biochemical characterization and structure determination. The receptor-bound pool consists of a filamentous conformation distinguishable from plaque Aß without any evidence for an oligomeric species.

## Receptor-bound pool of amyloid-ß

Our goal was to separate Aß species bound to receptor sites from free, unbound Aß and from plaque-associated fibrillary Aß. First, we processed fresh frozen autopsy brain tissue from cerebral cortex of Braak stage V/VI cases by Dounce homogenization in aqueous buffer followed by ultracentrifugation. A fraction containing monomers and free oligomers was released into the supernatant selectively from AD autopsy brain, but not neurologically intact control brain, and repeated extractions revealed near-complete depletion of this pool (Fig. 1a). Previously measured in vitro Aßo dissociation rates from receptors are a day or more at room temperature ^23^, so most receptor-bound Aß remained in the ultracentrifuge pellet together with Aß fibrils during these steps at 4C.

**Fig. 1.**
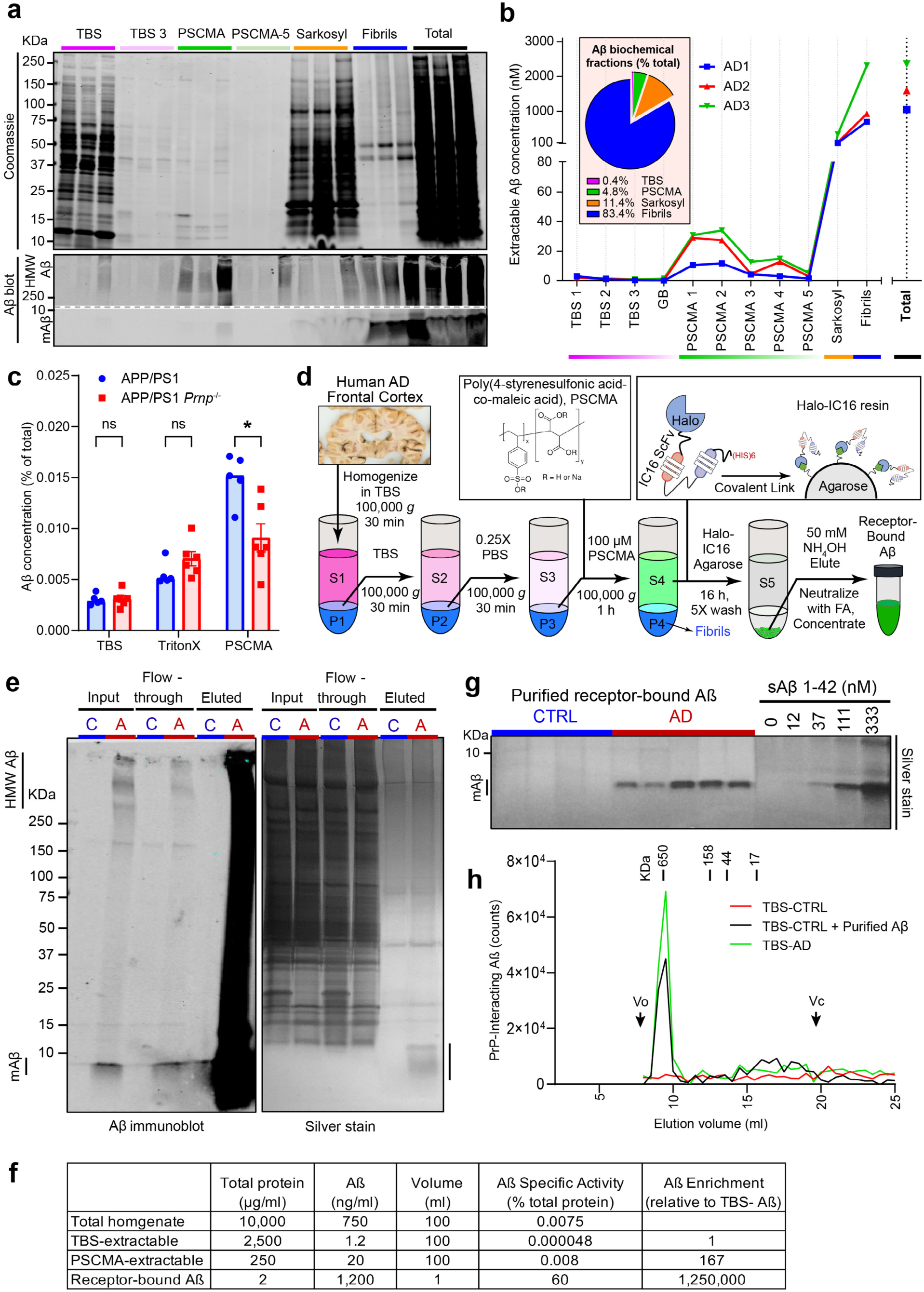
Purification of receptor-bound Aß. **a,** Coomassie total protein stain (top panel) and D54D2 anti-Aß immunoblot (bottom panels) of sequential biochemical protein extracts from 3 individual human AD brains. HMW and LMW regions of the same blot are displayed to show both SDS-resistant Aß assemblies and monomeric Aß. **b,** Quantitation of Aß in sequential biochemical extracts. Amount of Aß in sarkosyl-insoluble extracts, Aß fibrils and Total extractable Aß was measured by immunoblot, while all other (non-denaturing) fractions were measured by IC16/D54D2 ELISA. The pie chart insert represents the total extractable amount of Aß in each biochemical fraction as a percent of total brain Aß. **c,** Comparison of Aß levels in TBS, TBS-Triton (TBSX) and PSCMA sequential protein extracts from APP/PS1 transgenic AD mice with WT or Prnp^-/-^ background. Significantly lower amount of receptor-bound Aß is detected in APP/PS1, Prnp^-/-^ mice. Mean ± sem. Each dot is from a separate preparation from different mice. *, P < 0.05, one-way ANOVA with Tukey correction for multiple tests. **d,** Purification scheme allowing isolation of high purity native Aß from human AD brains. Brain homogenate is sequentially resuspended and centrifuged in TBS (twice), 0.25X PBS and PSCMA. The TBS soluble fractions are color-coded pink, PSCMA-extracted fractions are in green and insoluble pelleted material including fibrillar Aß is in blue. PSCMA-eluted receptor-bound Aß assemblies are immunopurified using IC16 ScFv covalently linked to Halo agarose. The inserts show the chemical structure of PSCMA and diagram of IC16-ScFv-HaloTag immobilization. **e,** D54D2 anti-Aß immunoblot (left) and total protein (Silver stain, right) of PSCMA extracted material (Input), immunodepleted PSCMA material (Flow-through) and IC16-immunopurified receptor-bound Aß (Elute) from control (C) and AD (A) brains. Strong increase in HMW Aß immunoreactivity is observed in eluted material, with >50% purity observed by silver stain. **f,** Aß purification table summarizing the total protein concentration, Aß content and Aß purity throughout the purification process. Total homogenate reflects all protein in brain tissue when extracted with directly 8M Urea with 4% SDS. **g,** Silver stain of purified receptor bound Aß from 5 different Alzheimer’s (AD) brains and comparable fraction from control (CTRL) brains. A dilution series of synthetic Aß42 standard was separated alongside brain-derived material on the same gel. No Aß is observed in any control samples while AD brains show a strong band at the expected Aß monomer mol wt. **h,** Size exclusion chromatography of purified receptor-bound Aß preparation. TBS lysates from control brain (TBS-CTRL), or control brain TBS lysate spiked with purified receptor-bound Aß from AD brain (TBS-CTRL + Purified Aß), or TBS lysate from AD brain (TBS-AD) were separated by size-exclusion chromatography on Superdex 200 column, and fractions assayed for PrP^C^-interacting Aß immunoreactivity (PLISA). Virtually no Aß immunoreactivity is observed in the control lysate, but addition of purified receptor-bound Aß results in a peak overlapping with the TBS-soluble Aßo peak observed in AD brain. Vo – void volume, Vc – column volume, numbers above the graph show the elution times of MW standards separated on the same column.

To release receptor-bound Aß species, we utilized the PrP^C^ antagonist, PSCMA, which was identified as a competitive blocker of Aß binding ^23^, and which is capable of dissociating human brain hydrogels composed of Aß and PrP^C^ receptors ^31^. Resuspension in the presence of micromolar concentrations of PSCMA released an Aß fraction containing ten-fold more total Aß immunoreactivity that was present in the sum of repeated TBS extracts from the same brain (Fig. 1b). This PSCMA-extracted material was a distinct pool, as demonstrated by sequential extractions yielding progressively less Aß release, and by the majority of the Aß species remaining in the ultracentrifuged pellet, as expected for Aß fibrils (Fig. 1b). Only a small amount of total protein was extracted by PSCMA in the receptor-bound Aß pool, providing a near 200-fold enrichment of Aß immunoreactivity per mg protein compared to the TBS-extractable pool (Fig. 1a, 1b).

Based on our previous work with recombinant Aß and purified PrP^C^, we hypothesized that the elution of receptor-bound Aß by PSCMA was largely the result of release from PrP-dependent hydrogels through a multivalent disruption of coacervation. We tested this hypothesis for the origin of receptor-bound Aß using a transgenic mouse model, APPswe/PS1ΔE9 (APP/PS1), on a wild-type versus *Prnp*^-/-^ background. The amount of Aß released from APP/PS1 brain was reduced by about 50% when PrP^C^ was absent (Fig. 1c). The PrP^C^ effect was specific to the receptor-bound Aß pool, since total Aß load and the free soluble Aß fractions were unchanged by *Prnp*^-/-^ status. The minor fraction of PSCMA-extractable Aß species in APP/PS1 brain lacking PrP^C^ may reflect the PSCMA sensitivity of alternate Aßo receptors.

The receptor-bound Aß species extracted by PSCMA was highly enriched, but not homogeneous. To further enrich the material, we developed a single chain anti-Aß antibody resin for moderate affinity-based purification, allowing removal of PSCMA and achieving rapid elution under mild conditions (Fig. 1d). Analysis of the final preparation by SDS-PAGE yielded a single prominent protein band by silver stain, which migrated at the same mobility as recombinant Aß42 peptide (Fig. 1e). There was no residual PrP^C^ by immunoblot analysis. Compared to TBS-soluble Aß immunoreactivity in human AD brain, the preparation was purified one million-fold (Fig. 1f). Similar purified receptor-bound Aß preparations were generated from cerebral cortex of five different Braak stage V/VI AD brain autopsy samples (Fig. 1g).

## Characterization of receptor-bound pool of amyloid-ß

We sought to assess immunologic and physical properties of the receptor-bound Aß pool from human AD brain. A panel of anti-Aß antibodies was utilized to compare the receptor-bound Aß from the four different AD brains to fibrillary Aß sedimenting in the presence of PSCMA from the same brains (Fig, 2a, 2b). For this comparison, the PSCMA-released Aß pool prior to antibody affinity retention was compared to PSCMA-resistant material containing plaque-associated Aß. The PSCMA-resistant material was further extracted with sarkosyl, and the sarkosyl-insoluble pellet resuspended for immunoblot. The Aß immunoreactivity with different amino-specific and carboxyl-specific antibodies was assessed after normalization to a mid-region anti-Aß antibody. The receptor-bound Aß pool was five-fold enriched for the most amino terminal residues of Aß1-42 (Fig, 2a, 2b). The level of reactivity for pyroglutamylated Aß was similar in the two preparations. At the carboxyl terminus, reactivity was similar for the two pools with anti-Aß42 and anti-Aß43 specific antibodies, but greater for Aß40 in the receptor-bound pool. These differences confirm the distinct nature of the receptor-bound Aß species relative to fibrillary Aß and show more extensive proteolysis of the amino-terminus for the insoluble fibrillary species. The FDA approved anti-Aß antibody, Lecanemab, is reported to have preferential affinity for oligomeric and protofibrillary Aß species, with less avidity for Aß filaments and minimal interaction with monomeric Aß ^32^. We confirmed its high affinity for the receptor-bound Aß species (Extended Data Fig. 1).

Next, we characterized the physical size of the receptor-bound Aß pool by electron microscopy (EM) and size exclusion chromatography (SEC). Receptor-bound Aß exhibited a rod-like shape in EM images with a length of 60±23.4 nm, 70±24.4 nm and 71±29.2 nm (mean ± sd) from three different AD brains (Fig. 2c), demonstrating a filamentous nature for the receptor-bound Aß assemblies in multiple brain samples. To characterize size using SEC, the purified receptor-bound Aß needed to be first mixed with a protein-rich preparation to avoid non-specific adsorption during chromatography. SEC fractions of TBS extracts from control brain are protein-rich but show no background of detectable PrP^C^-interacting Aß immunoreactivity (Fig. 1h). When these TBS extracts are mixed with purified receptor-bound Aß from AD brain, there is a single PrP^C^-interacting Aß immunoreactivity peak with a size matching migration of globular proteins in the 650 kDa range, consistent with the EM appearance and an assembly of about 250 Aß monomers (Fig. 1h). This is similar to the SEC elution profile for free TBS-soluble Aßo species from AD brain.

**Fig. 2.**
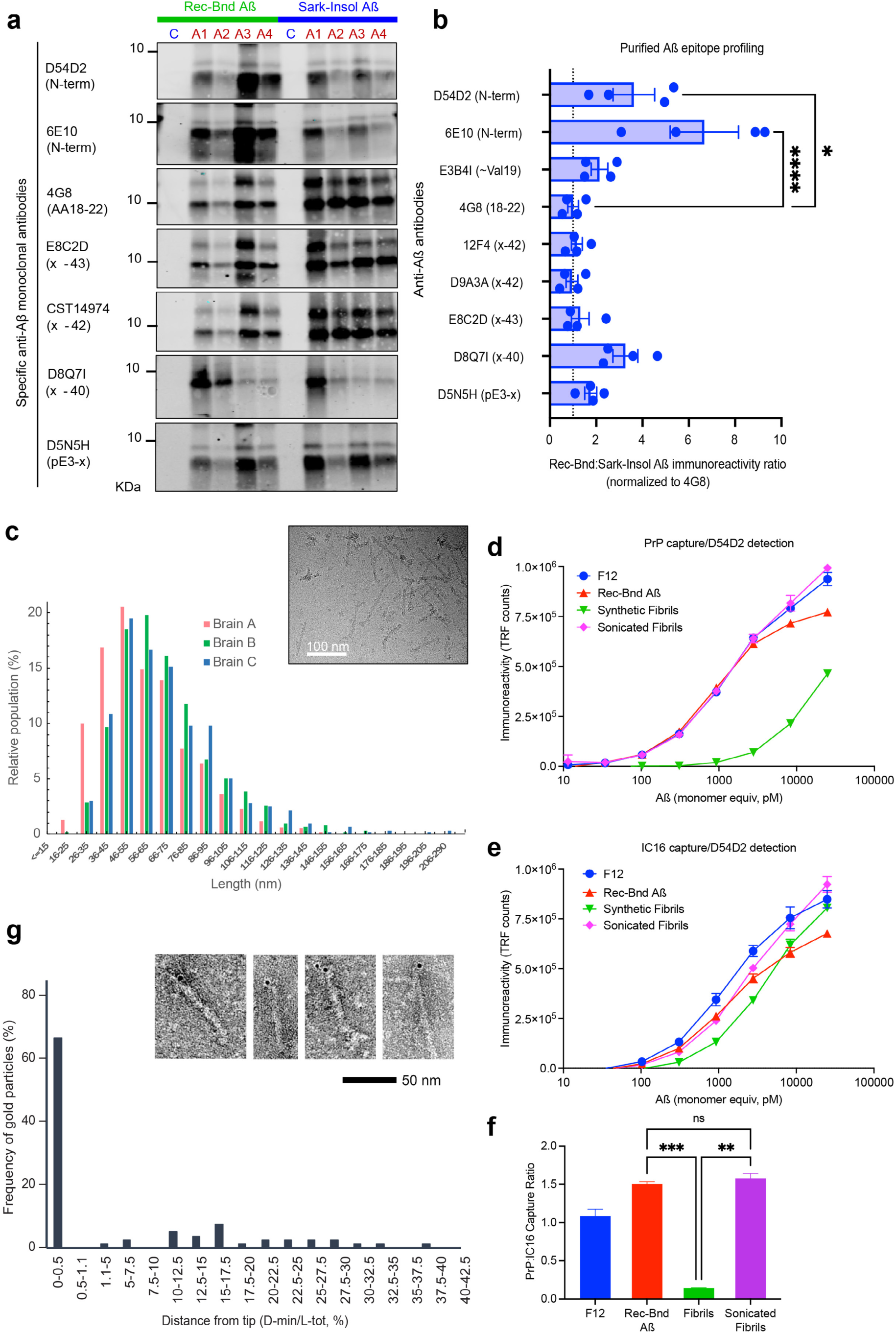
Biochemical characterization of receptor-bound Aß. **a,** Immunoblot epitope profiling of HFIP-monomerized purified receptor-bound Aß (Rec-Bnd Aß) compared to sarkosyl-insoluble Aß fibrils extracted from the same brain (Sark-Insol Aß). One control brain (C) and 4 individual AD brains (A1-A4) were used. Epitope location within Aß sequence is indicated next to the antibody clone name on the left. **b,** Graph of quantified immunoreactivity in A. Signal is normalized to 4G8 antibody (mid-section of Aß, AA18-22). The ratio of Receptor-bound to Sarkosyl-insoluble Aß immunoreactivity is plotted. Note significant increase in the immunoreactivity with antibodies directed against N-terminus of Aß in the PSCMA fraction. Each dot is the determination from a different autopsy brain. Mean ± sem for n=4. One-way ANOVA for each antibody versus 4G8, with Dunnett correction. * P<0.05, **** P<0.0001. **c,** Length distribution of purified Aß assemblies determined by transmission electron microscopy. The inset shows a representative image of Aß assemblies using cryo-EM. **d,** PLISA characterization of various Aß preparations. Alzheimer’s brain-derived receptor-bound Aß binds PrP^C^ with the affinity similar to that for synthetic Aß oligomeric preps (F12), while synthetic fibrils have much lower affinity (Aß monomer equivalent concentrations). However, sonicating the fibrils increases their PrP^C^ binding. Mean ± SD for n=3. **e,** IC16 capture antibody, D84D2 detection antibody ELISA characterization of various Aß preparations. Similar affinities observed for both fibrillar and oligomeric synthetic Aß preparations, with exception of globulomer Aßo having much higher activity. Mean ± SD for n=3. **f,** The ratio of the PrP^C^ to IC16 capture signal from the assays in d, e is plotted for data from the 936 pM Aß monomer equivalent value. Values are normalized to the ratio for F12 Aßo preparation. Mean ± SD for n=3. Welch’s ANOVA, with Dunnett correction. **P<0.01, ***P<0.001. **g,** Localization of PrP^C^ binding sites to receptor-bound Aß detected by immunogold labeling as a function of distance from the tip to the middle of the rod. The inset shows representative images of brain-derived Aß assemblies using negative stain EM after incubation with purified PrP^C^, mouse anti-PrP^C^ antibody and colloidal gold labeled anti-mouse secondary antibody.

## Signaling by receptor-bound pool of amyloid-ß

As an Aß receptor, PrP^C^ is of high affinity and essential for a range of phenotypes in AD models. We compared the PrP^C^ binding properties of the human brain receptor-bound Aß pool with that of other Aß preparations (Fig. 2d). Expressed in monomer equivalents, the apparent avidity in a plate-based PrP^C^-interacting Aß immunoreactivity (PLISA) assay was similar to that for recombinant Aß42 globulomer (12mer), a synthetic Aß42 ADDL preparation in F12 medium (apparent M_R_ of 500,000), as well as unpurified TBS-soluble free Aßo from human AD brain (apparent M_R_ of 1,000,000). In contrast, the affinity of Aß42 monomers or synthetic Aß42 fibrils or sarkosyl-insoluble Aß fibrils from human brain for PrP^C^ was nil or at least two orders of magnitude lower.

The interaction of rod-like shape of human brain receptor-bound Aß with PrP^C^ raises the question of whether receptor interactions occur along the side, or at the tips, two clearly different environments. Since the principal EM appearance difference between receptor-bound Aß and insoluble fibrillary Aß is a higher ratio of tip to shaft length, tips may be key for PrP^C^ interaction. We compared PrP^C^ binding of intact Aß42 synthetic fibrils to sonicated preparations that had been sheared to a length of 50-100 nm as confirmed by negative stain transmission EM. The breakage of fibrils and the creation of multiple tips supported PrP^C^ binding to the otherwise non-interacting Aß42 fibril preparation, with an increase in avidity of greater than 20-fold (Fig. 2d). This strongly suggests that tip concentration is crucial for PrP^C^ interaction. In contrast to PrP^C^ binding, detection of oligomeric Aß species using two competing antibodies was not altered by sonication of fibrils to smaller size (Fig. 2e). The ratio of Aß capture by PrP^C^ versus antibody from the fibril preparation increased after fragmentation to levels matching that of receptor-bound Aß (Fig. 2f).

To observe interaction of Aß filament tip with PrP^C^ directly, we incubated human brain receptor-bound Aß with recombinant PrP^C^, applied to an EM grid and stained with anti-PrP antibody (3F4) and immunogold anti-mouse antibody. Gold particles associated with Aß structures were localized relative to the tip position, and nearly all particles were within 5 nm of the tips (Fig. 2g). A similar conclusion has been suggested from light microscopy studies of fluorescently tagged synthetic Aß42 peptide ^33^. This confirms discrete localization of binding sites for this receptor and explaining the strong binding preference of PrP^C^ for oligomeric Aß over intact Aß fibrils.

Since PrP^C^ binds to the tips of receptor-bound Aß, its presence is predicted to inhibit further polymerization in the presence of Aß42 monomers. Indeed, the presence of recombinant PrP^C^ reduced the rate of Aß42 polymerization or Aß42 polymerization seeded by fibrils (Extended Data Fig. 2). This finding is consistent with several *in vitro* kinetic reports ^34^, although PrP^C^ deletion does not modify the extent of chronic *in vivo* Aß plaque accumulation in mouse models ^21, 25, 30^ where multiple metabolic factors are at play.

Various preparations of Aßo from synthetic, recombinant or brain sources have been shown to alter synaptic signaling acutely and induce anatomical synapse loss, both of which can be blocked by PSCMA ^23^. We assessed biological activity by exposure of human induced pluripotent stem cells (iPSC)-derived cortical excitatory neurons (60 days post differentiation by Neurogenin2 induction) to the purified receptor-bound Aß species (16-160 pM assembly, 4-40 nM monomer equivalent) for 7 days. The density of synaptic profiles identified by overlap of Synapsin1 and Homer1b/c staining at MAP2-immunoreactive dendrites was significantly reduced by receptor-bound Aß from three different AD brains as compared to control brain samples (Extended Data Fig. 3). These results confirm a potent biological effect of receptor-bound Aß, similar to that described previously for other species.

## Cryo-EM of receptor-bound amyloid-ß

We utilized cryo-EM to determine the structure of receptor-bound Aß purified from three AD brains. In each case, the majority of images yielded an electron density map that we term Conformation 1 of the receptor-bound Aß with a resolution of 2.8Å from Brain A (Fig. 3a, 3d, 3f, Extended Data Fig. 4). A less prevalent Conformation 2 with resolution of 3.1Å from Brain A and Brain B was also observed (Fig. 3b, 3e). Atomic models were built from these maps and included Aß42 residues 9-42 (Fig. 4a-d). These models are composed of two symmetric folded S-shaped Aß peptides in each rung of a twisting amyloid filament. The rungs are spaced at 4.75 Å, so a 65 nM length implies a total of about 300 monomers per assembly (Fig. 4a, 4c). Residues 1-8 are not resolved but may account for fuzzy electron density extending from residue 9 and lying alongside the filament.

**Fig. 3.**
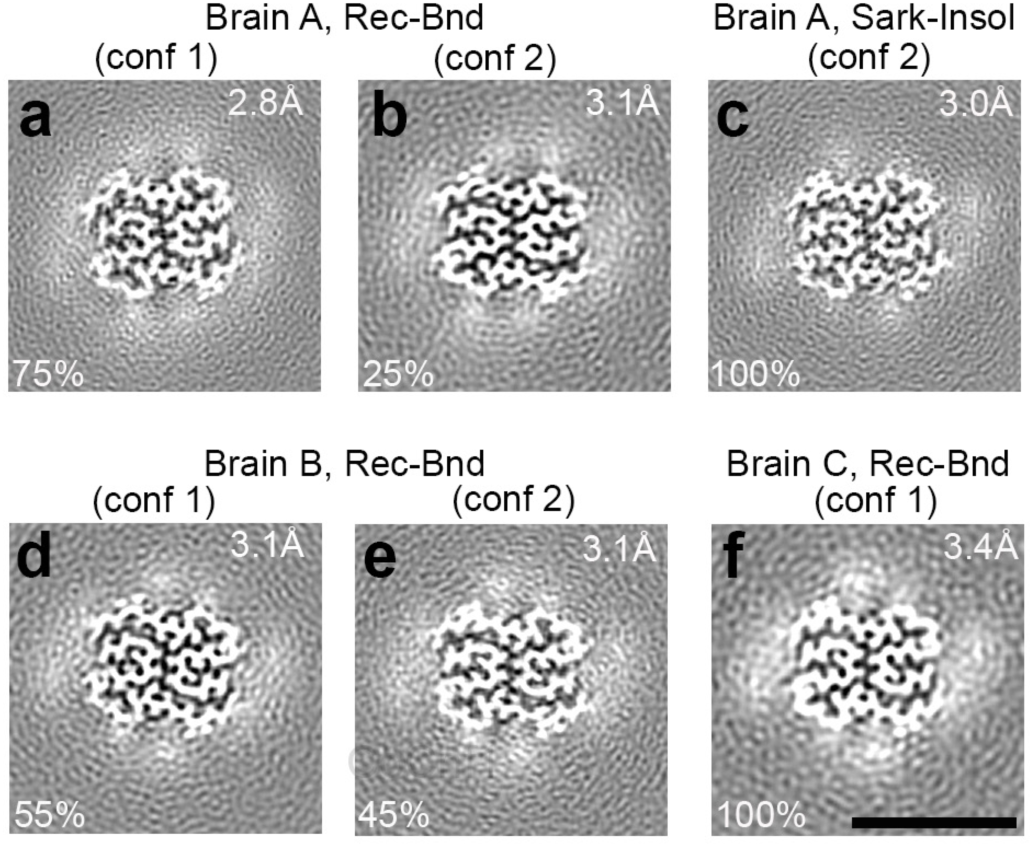
Three-dimensional reconstruction of receptor-bound Aß purified from Alzheimer’s brain. **a, b, d-f,** Central cross-sections from the experimental electron density maps of the PSCMA-extractable receptor-bound Aß in a plane perpendicular to the fibril axes. The projected thickness is approximately one Aß-rung. High-density is white. Two similar but different conformations (labeled as Conformation 1 and Conformation 2) are revealed in the cross-sections. Percentages of conformations are shown in the bottom-left corner. The resolutions of the cryo-EM maps are shown in the upper-right corner. **c,** Central cross-section from the experimental electron density maps of the sarkosyl-insoluble Aß amyloid fibrils. The postmortem brain tissues of Alzheimer’s patients are labeled Brain A, Brain B, Brain C. Bar, 5 nm.

**Fig. 4.**
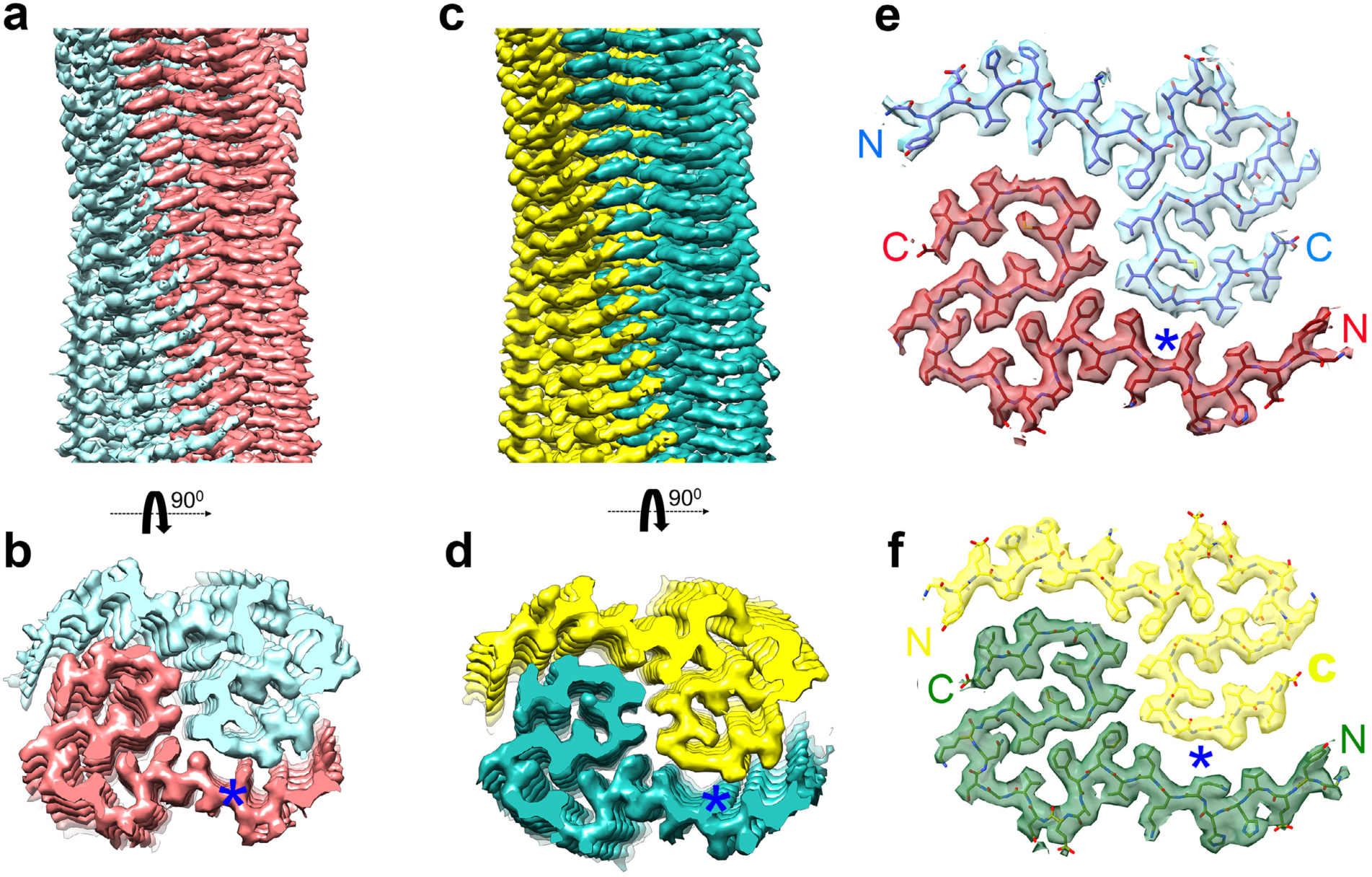
Surface renderings of receptor-bound Aß cryo-EM reconstructions. **a, b,** Surface renderings of the cryo-EM reconstruction showing a side view (**a**) and top view (**b**) of the receptor-bound Aß in Conformation 1 from Brain A. Coloring (in red and light cyan) is by Aß42 peptide monomer taking into account the helical and pseudo-two-fold symmetry. **c, d,** Surface renderings of the cryo-EM reconstruction showing a side view (**c**) and top view (**d**) of the receptor-bound Aß in Conformation 2 from Brain B. Coloring (in green and yellow) is by Aß42 peptide monomer taking into account the helical and pseudo-two-fold symmetry. **e,** Cross-sectional view shows the model of two Aß42 peptides built in the electron density of one rung from Conformation 1 (Brain A). Coloring (in red and light cyan) is by Aß42 peptide monomer. The N- and C- terminal ends are highlighted. **f,** Cross-sectional view shows the model of two Aß42 peptides built in the electron density of one rung from Conformation 2 (Brain B). Coloring is by Aß42 peptide monomer. The N- and C- terminal ends are highlighted. Blue asterisks highlights Q15 difference between the two conformations.

The two Conformations differ from one another in the amino terminal region with a different rotameric state for the sidechain of Q15 (Fig. 4b, 4e in comparison to 4c, 4f), and a shift in the peptide backbone (Fig. 5a, 5b, 5c). This results in a different pattern of intra-molecular hydrogen bonding for V12-Q15 (Fig 5a, and 5d, 5e versus 5g, 5h). In Conformation 1 there is 1:1 bonding to the next rung in the filament at each of these amino acid residues. For example, the amide amino group Q15 sidechain to the amide carbonyl of Q15 sidechain in the next rung (Fig. 5d, 5e). In contrast, Conformation 2 bonding to the next rung is more frequently shared by two residues in the next rung, for example the amide amino group Q15 sidechain to both carbonyl of V12 backbone and amide carbonyl of Q15 sidechain in the adjacent rung (Fig. 5g, 5h).

**Fig. 5.**
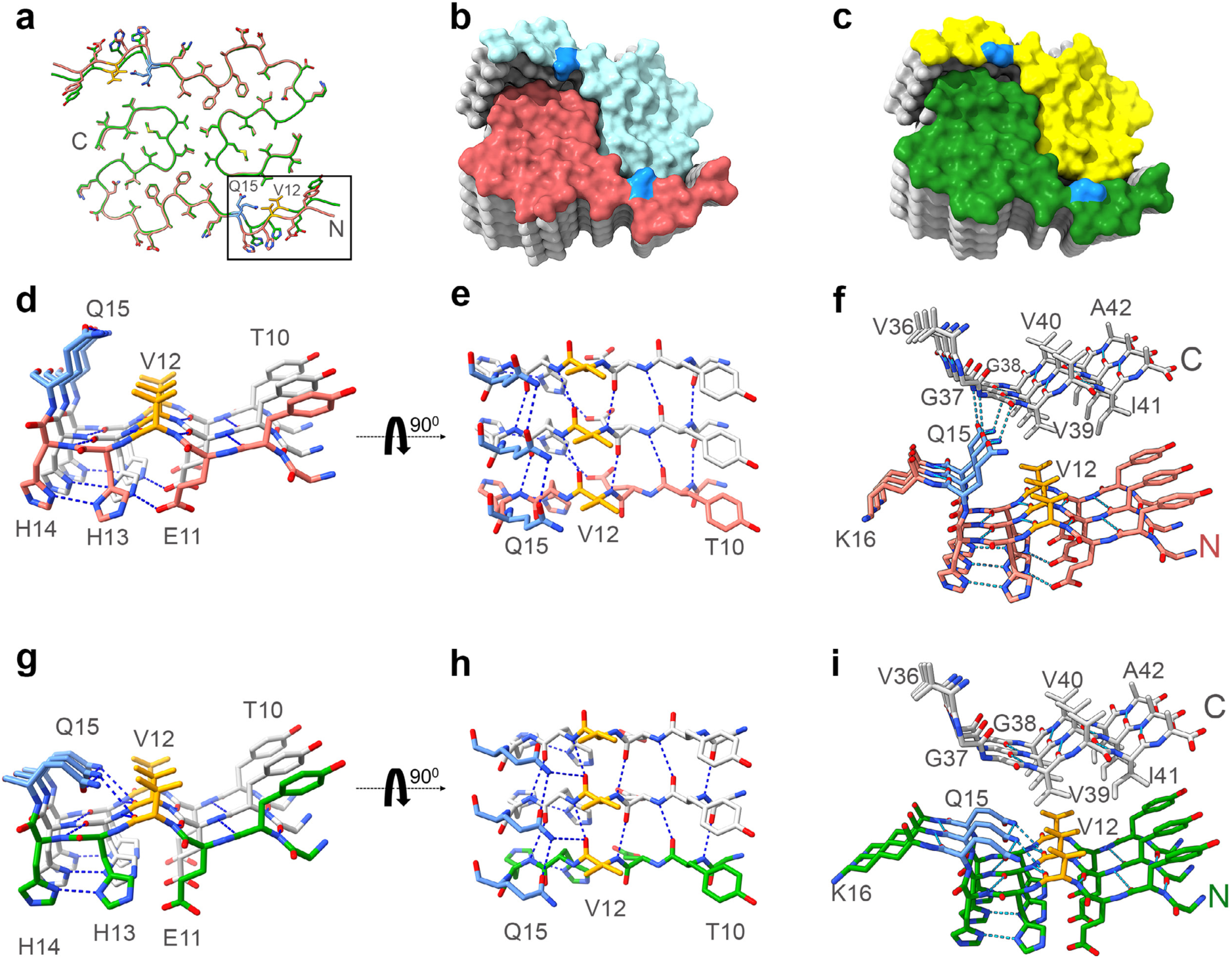
Comparison of two conformations adopted by receptor-bound Aß. **a,** Overlay of the molecular models of one Aß rung from Conformation 1 (Brain A, in red) and of one Aß rung from Conformation 2 (Brain B, in green). The rectangle highlights the main differences between the two conformations. Side chain residues are shown in stick representation. Key side chains Q15 (colored in blue) and V12 (colored in orange) are highlighted. N- and C- mark the amino and carboxyl ends of an Aß42 peptide monomer. **b,** Surface representation of the packing assembly of 5 rungs with the Aß42 peptide monomer in Conformation 1 (resolved from Brain A). The rung at the top is colored by the peptide monomer in red and cyan, respectively. The location of the Q15 positions on the surface of the top rung is illustrated in blue. **c,** Surface representation of the packing assembly of 5 rungs with the Aß42 peptide monomer in Conformation 2 (resolved from Brain B). The rung at the top is colored by the peptide monomer in green and yellow, respectively. Q15 locations on the surface of the top rung are illustrated in blue. **d, e,** A close-up view of the region highlighted with a rectangle in panel (**a**) for residues T10-Q15 from 3 rungs of the Aß peptide in Conformation 1 (resolved from Brain A). Hydrogen-bonding pattern of backbone atoms within the intermolecular parallel β-sheet is shown in dotted lines. Point of view is rotated 90 degrees between **d** and **e**. **f,** Residues V36-A42 of Aß42 peptide from 3 rungs of the opposing protofilament (colored in gray) are shown together with the region from **d, e** for Conformation 1 (colored in red). Hydrogen-bonding pattern of the Q15 sidechain is shown with dotted lines contacting the opposing protofilament peptides. **g, h,** A close-up view of the region highlighted with a rectangle in panel (**a**) for residues T10-Q15 from 3 rungs of the Aß42 peptide in Conformation 2 (resolved from Brain A). Hydrogen-bonding pattern of intramolecular and intermolecular side chains and backbone atoms is shown in dotted lines. Point of view is rotated 90 degrees between **g** and **h**. **i,** Residues V36-A42 of Aß42 peptide from 3 rungs of the opposing protofilament (colored in gray) are shown together with the region from **g, h** for Conformation 2 (colored in green). Hydrogen-bonding pattern of the Q15 sidechain is shown with dotted lines, and there are no contacts with the opposing protofilament Aß peptide.

The different rotameric state of the sidechain of Q15 also affects the intermolecular interactions between Aß peptides in the two opposing protofilaments (Fig. 5f versus 5i). In Conformation 1, the amide group of Q15 sidechain within one Aß peptide forms a hydrogen bond with both the G38 backbone carboxyl and amino terminus from the opposing Aß peptide in an adjacent rung (Fig. 5f). These interactions may stabilize the greater twist observed for Conformation 1. In comparison, the orientation of Q15 in Conformation 2 does not allow for such interactions (Fig. 5i).

Accompanying this monomer change, the stacking and twist of monomers differs between Conformation 1 and Conformation 2 (Fig. 4a and 5b for Conformation 1 versus 4c and 5c for Conformation 2). The angle between the plane of one peptide and its partner in the same rung is 178.16 degrees for Conformation 1 and 178.52 degrees for Conformation 2 (Table 1). Over the 65-nm length of the receptor-bound Aß42 rod containing about 300 monomers, the position of the Aß42 monomer rotates 507 degrees around the long axis in Conformation 1 versus 407 degrees for Conformation 2. Thus, the two ends of a 65 nm rod differ by 100 degrees of rotation. More strikingly, amino terminal residues 9-15 of one Aß42 rung are adjacent to carboxyl residues of a monomer two rungs below in Conformation 1, but one rung below in Conformation 2 (compare Fig. 5b and 5c; Extended Data Fig 5).

**Table 1.** Cryo-EM Data Collection and Processing.

|  | Brain A<br>(Receptor-Bound) |  | Brain B<br>(Receptor-Bound) |  | Brain C<br>(Receptor-Bound) |  | Brain A<br>(sarkosyl-insoluble) |
| --- | --- | --- | --- | --- | --- | --- | --- |
| Data Collection |  |  |  |  |  |  |  |
| Microscope | Titan Krios G2 |  |  |  |  |  |  |
| Voltage (kV) | 300 |  |  |  |  |  |  |
| Magnification | 81,000 x |  |  |  |  |  |  |
| Pixel size (Å/pixel) | 1.068 |  |  |  |  |  |  |
| Camera | Gatan K3 with EF |  |  |  |  |  |  |
| Exposure rate (e-/px/s) | 21.5 |  | 16.1 |  | 16.1 |  | 25.9 |
| Exposure time | 2.155 |  | 2.936 |  | 2.936 |  | 1.846 |
| Number of frames | 39 |  | 42 |  | 42 |  | 42 |
| Electron Dose (e-/Å²) | 41 |  | 41 |  | 41 |  | 42 |
| Defocus range (μm) | -2.5 to -1.25 |  | -2.5 to -1.25 |  | -2.5 to -1.25 |  | -2.5 to -1.25 |
| Data processing |  |  |  |  |  |  |  |
| Number of movies | 2,464 |  | 2,285 |  | 1,238 |  | 2,493 |
| Extraction box size (pixels) | 200 |  | 200 |  | 200 |  | 200 |
| Initial number of particle projections | 604,506 |  | 247,940 |  | 212,112 |  | 575,508 |
|  | Conf1 | Conf2 | Conf1 | Conf2 | Conf1 |  | Conf2 |
| Final number of particle projections | 160,787 | 51,098 | 62,771 | 52,021 | 11,500 |  | 80,229 |
| Symmetry imposed | C1 |  | C1 |  | C1 |  | C1 |
| Map resolution in Å (FSC 0.143) | 2.8 | 3.1 | 3.1 | 3.1 | 3.4 |  | 3 |
| Map sharpening B factor (Å²) | -91.68 | -95.08 | -103.69 | -91.68 | -83.43 |  | -82.04 |
| Helical rise (Å) | 2.355 | 2.37 | 2.345 | 2.36 | 2.354 |  | 2.37 |
| Helical twist (°) | 178.164 | 178.52 | 178.168 | 178.5 | 178.144 |  | 178.5 |
| Structure refinement |  |  |  |  |  |  |  |
| Initial model used | 7Q4B |  | 7Q4B |  |  |  | 7Q4B |
| Model composition |  |  |  |  |  |  |  |
| Protein residues | 340 |  |  | 340 |  |  | 340 |
| R.m.s. deviations |  |  |  |  |  |  |  |
| Bond lengths (Å) | 0.009 |  |  | 0.004 |  |  | 0.006 |
| Bond angles (°) | 0.687 |  |  | 0.464 |  |  | 0.486 |
| Validation |  |  |  |  |  |  |  |
| MolProbity score | 0.64 |  |  | 1.23 |  |  | 0.58 |
| Clashscore | 0.4 |  |  | 3 |  |  | 0.2 |
| Poor rotamers (%) | 0 |  |  | 0 |  |  | 0 |
| Ramachandran plot |  |  |  |  |  |  |  |
| Favored (%) | 100 |  |  | 96.88 |  |  | 100 |
| Allowed (%) | 0 |  |  | 3.12 |  |  | 0 |
| Disallowed (%) | 0 |  |  | 0 |  |  | 0 |

We considered whether each receptor-bound filament contained only one Conformation of Aß or the two Conformations were intermixed in a single filament. The positions of filament segments contributing to the resolved conformations were traced on individual filaments in the cryo-EM micrographs. This shows that certain filaments contained distinct segments composed of Conformation 1 and Conformation 2, while other filaments contained only Conformation 1 and less frequently only Conformation 2 (Extended Data Fig. 6). Thus, Conformation 1 and Conformation 2 can exist within the same filament and the shift in residues V12-Q15 does not require filament breakage.

A small minority of receptor-bound Aß filaments exhibited a greater diameter and electron density maps revealed a doublet of the two symmetric folded S-shaped Aß peptides observed in the more prevalent Conformations (Extended Data Fig. 7). Although the limited number of these doublet filaments prevented 3D reconstructions with high resolution, it was clear that the doublets from Brain A and Brain B were composed of Aß in Conformation 2, as opposed to Conformation 1, from the observation of density corresponding to Q15. The twist of Conformation 1 may restrict the formation of doublets.

The structure of the two receptor-bound Aß species was compared to reported Aß structures isolated from Alzheimer’s brain. The less prevalent Conformation 2 is closely similar to the filament structure reported for sarkosyl-insoluble plaque material throughout Aß9-42 (Extended Data Fig. 8) ^10^. In contrast, Conformation 1 shows the same degree of deviation from sarkosyl-insoluble Aß filaments as it does from Conformation 2. We considered whether this might be due to subject-specific conformations, or due to different conformations within the same brain for receptor-bound Aß as compared to filamentous PSCMA-resistant plaque-associated Aß. We therefore examined material from Brain A which remained in the ultracentrifuge pellet after sequential extraction with TBS, PSCMA and sarkosyl. The resulting sarkosyl-insoluble material had at least three filamentous forms visible by cryo-EM (Extended Data Fig. 9). One type showed electron density maps consistent with Aß filaments, one with Tau filaments and one with TMEM106B filaments. The electron density map for Aß filaments resolved at 3.0 Å (Fig. 3c), and an atomic model matched the published sporadic Alzheimer Aß42 filament and Conformation 2 very closely (0.265Å rmsd and 0.160 Å rmsd, respectively), while differing from receptor-bound Aß Conformation 1 from the same brain (1.503 Å rmsd) (Fig. 6). Conformation 1 is most similar to a lower resolution structure isolated from Down’s syndrome brain ^35^. These data support the enrichment of distinct Aß42 conformations in these different pools of Aß.

**Fig. 6.**
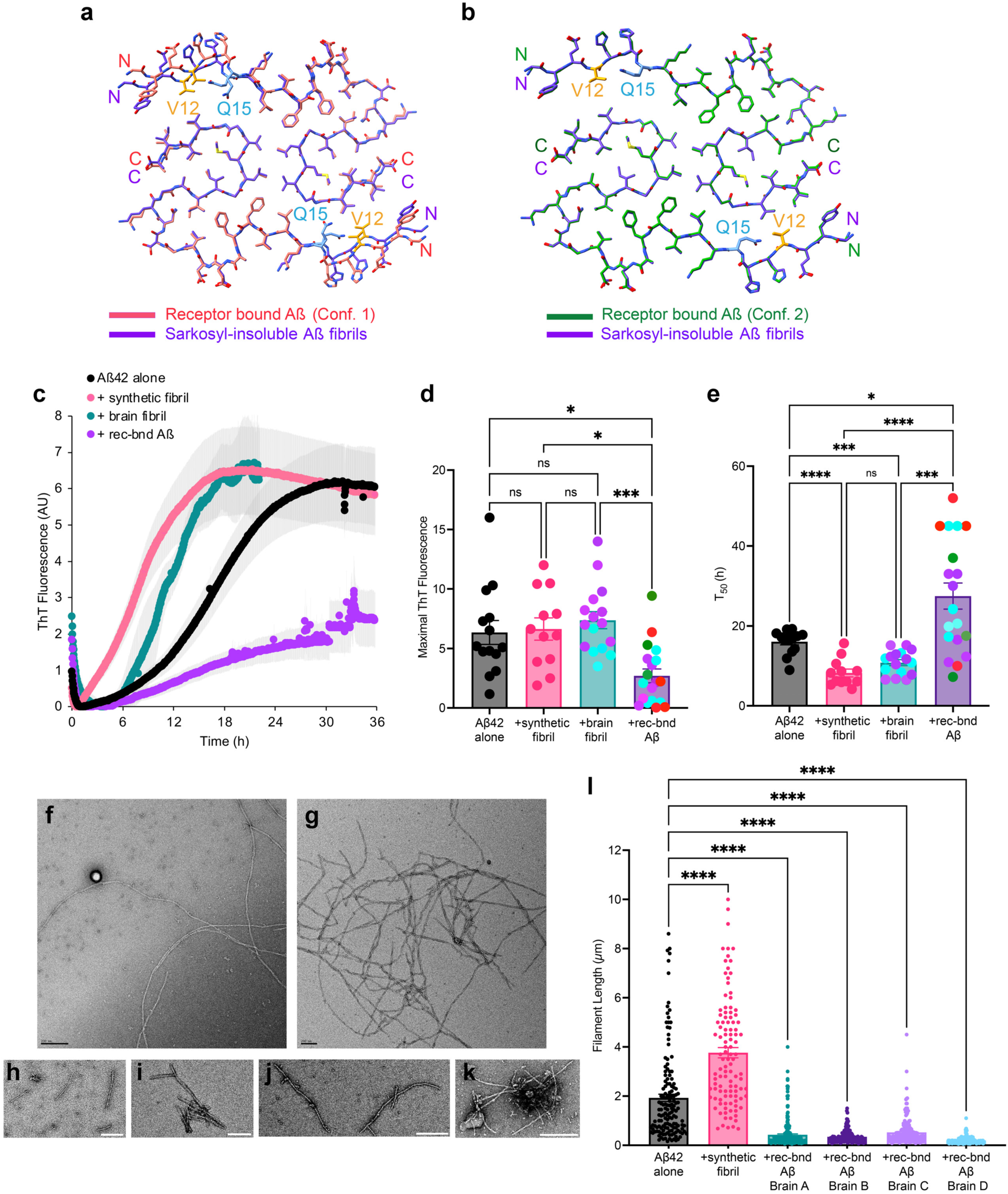
Differential seeding of Aß42 polymerization by receptor-bound Aß and sarkosyl-insoluble Aß fibrils. **a,** Overlay of the molecular models of one rung from Conformation 1 (Brain A, receptor-bound Aß in red, PSCMA) and of one rung of the Aß42 filaments found in the sarkosyl-insoluble preparations of Brain A (in purple). Atoms are shown in stick representation. r.m.s.d over 68 C_α_ atoms 1.505 Å. **b,** Overlay of the molecular models of one rung from Conformation 2 (Brain A, receptor-bound Aß oligomer in green, PSCMA) and of one ß-rung of the Aß42 filaments found in the sarkosyl-insoluble preparations of Brain A (in purple). Atoms are shown in stick representation. r.m.s.d over 68 C_α_ atoms 0.160 Å. **c,** Amyloid seeding assays monitored by Thioflavin T fluorescence over 36 hours. Reactions include 5 µM synthetic Aß42 monomer alone (n=12 experiments each), or with addition of synthetic Aß42 fibril seeds (20 nM monomer equivalent, n=12 experiments), or with addition of brain sarkosyl-insoluble Aβ42 fibril seeds (brain fibril, 20 nM monomer equivalent, n=5 experiments with Brains A-B), or with addition of purified receptor-bound Aß seeds (20 nM monomer equivalent, n=19 experiments with Brains A-D). Data are mean ± SEM. **d, e,** The maximal fluorescence ThT fluorescence (**d**) and the time to reach half maximal ThT fluorescence (**e**) for each Aß42 polymerization assay in C is presented. Results for sarkosyl-insoluble Aß42 fibril seeds (brain fibril) from Brain A and B, and purified receptor-bound Aß seeds from Brains A-D are marked with different color dots. Data are mean ± SEM. Welch ANOVA tests with Dunnett correction for multiple tests. *P<0.05, **P<0.01, ***P<0.001. **f-k,** Representative negative staining electron microscopy of filament assembly at the end of the polymerization reactions. Examples are shown for long filaments from 5 µM synthetic Aß42 alone (**f**) and from seeding reactions of 5 µM synthetic Aß42 monomer with 20 nM of synthetic Aß42 fibril seeds (**g**) as well as shorter filaments from seeding reactions of 5 μM synthetic Aß42 monomer with 20 nM of receptor-bound Aß from Brain A (**h**), Brain B (**i**), Brain C (**j**) and Brain D (**k**). Scale bar, 200 nm (**f, g, h, j, k**). Scale bar, 100 nm (**i**). **l,** Filament length measurements from the negative-staining micrographs of the polymerization reactions as in **f-k**. Mean ± SEM. One-way ANOVA tests with Dunnett correction for multiple tests. *P<0.05, **P<0.01, ***P<0.001.

## Impact of receptor-bound Aß conformation

The biochemical separation and structural analysis of receptor-bound Aß reveals a different filament conformation from that of filamentous plaque Aß. We considered whether the two conformations might exhibit different fibril-seeding properties when compared directly under similar conditions. We seeded the two AD brain species at 20 nM monomer equivalent concentration into solutions of 5 µM monomeric Aß42 (1:250 molar ratio) and monitored polymerization rates using thioflavin T fluorescence (Fig. 6c-h). We also assessed Aß fibrils prepared from synthetic peptide *in vitro* and similarly fragmented by sonication. As expected, the seeding rate of Aß42 was accelerated about two-fold in the presence of seeds from either sarkosyl-insoluble Aß or synthetic fibrils as compared to Aß42 monomers without seeds (Fig. 6d). The maximal fluorescence was similar with and without fibril seeds (Fig. 6e). In contrast, the receptor-bound Aß species significantly slowed polymerization and reduced the maximal fluorescence achieved (Fig. 6c-e). We examined the material present at the completion of the seeding reaction by negative stain EM (Fig. 6f,g). The amyloid filaments generated with no seeds, sarkosyl-insoluble Aß seeds or synthetic fibril seeds were long, averaging more than 2,000 nm in length. The filament lengths generated by seeding with receptor-bound Aß purified from 4 different brains were significantly shorter, with most in the 200-500 nm length range. These data are consistent with the receptor-bound Aß seeding a different conformation than does the filamentous plaque Aß or synthetic Aß fibrils.

## Discussion

We have isolated, purified to homogeneity, characterized and determined the structure of the receptor-bound Aß pool from sporadic Alzheimer’s disease brain. The receptor-bound pool is ten-fold more prevalent than the free soluble Aß fraction. Receptor-bound Aß is uniform in shape, being composed of about 65 nm long rods containing on the order of 300 monomers in an amyloid filament. The distinctness of this pool is demonstrated by sequential extractions, by PrP^C^ dependence, by cryo-EM-determined structure, and by seeding properties. By virtue of its elution from brain receptor sites, the receptor-bound Aß species has the most direct relevance for disease pathophysiology.

These studies of receptor-bound Aß fail to reveal an “oligomer” bound to receptors. Previous work had postulated an oligomeric state based on dimer or trimer assemblies, but a defined oligomer has never been isolated and characterized at the atomic level. Moreover, one original report supporting an oligomeric form has been retracted^36^. Based on our observations of receptor-bound Aß here, the participation of an oligomeric state for Aß unrelated to amyloid filaments in Alzheimer’s pathophysiology is unlikely.

While the structure of receptor-bound Aß substantially overlaps with Aß filaments from insoluble fractions that are largely plaque-derived, the cryo-EM-derived atomic structure of the two forms are distinct, even when comparing fractions from a single brain. The altered conformation of amino terminal Aß residues 9-15 is coupled with different tilts for Aß monomers in each rung, and a different twist for the receptor-bound Aß filament. Receptor-bound Aß abundance may be crucial for disease activity and progression. In particular, the concentration of Aß assembly tip ends and the presence of Conformation 1 may be key factors for neuronal damage in Alzheimer’s. The temporal relationship between receptor-bound Aß and fibrillary Aß is not yet clear. They may interconvert freely or sequentially or be relatively separate non-interchangeable pathways of polymerization. However, the different states of irreversible proteolytic processing implies that the pools are not fully interconvertible within the time course of Aß proteolysis.

PrP^C^ binds several different ß-sheet rich Aß assemblies, including globulomers, ADDLs, TBS-soluble human brain Aßo, and receptor-bound Aß. While fibrillary Aß from synthetic or human brain sources shows little affinity for PrP^C^, fragmentation to create filament tips supports robust PrP^C^ avidity. These findings imply a specific interaction of PrP^C^ with filament tips and a role in titrating Aß filament elongation but not side-to-side templating. While PrP^C^ binds to tip ends of different conformations, it may favor or induce the more twisted Conformation 1 by virtue of its presence. Regardless, this observation highlights the potential importance of assessing tip end concentration in biological samples.

Why are the receptor-bound Aß limited to the 65 nm size range? This may reflect an equilibrium between the rate of Aß polymer extension at brain Aß monomer concentrations and the capping of Aß filament ends determined by brain concentrations of receptor binding sites such as PrP^C^. Alternatively, the twist of rungs and the tilting of monomers within rungs for Conformation 1 may make further polymerization unfavored. Our comparative seeding assays support the hypothesis that the receptor-bound Aß conformation is a distinct “strain” of Aß amyloid.

The structure of free unbound, soluble Aßo is not explored here. A recent study in which chunks of minced brain tissue were soaked to release these species has suggested that they may be most similar to insoluble Aß filaments of Conformation 2 ^16^.

Previous studies have indicated different conformations for Aß filaments from inherited Alzheimer’s cases and from mouse transgenic preparations ^10, 12^. We have not assessed receptor-bound Aß species from these sources. Studies of these alternate sources may further illuminate the relationship between insoluble fibrillary Aß and receptor-bound Aß. If receptor-bound conformations are shared between sources while fibrillary conformations are not, this would support a hypothesis that the Conformation 1 and Conformation 2 are not exchangeable and are distinct polymerization paths.

Therapeutic anti-Aß antibodies have recently received regulatory approval. Of note, Lecanemab development included a focus on the APP arctic mutation as an antigen which favors protofilament and oligomer reactivity ^3, 32^, and we show that receptor-bound Aß is recognized by this antibody. Donanemab development was based on pyroglutamylated forms of Aß ^4, 37^, which we show is present equally in both receptor-bound Aß and insoluble fibrillary Aß. The efficacy of these antibodies may be derived from clearance of the receptor-bound Aß conformation described here. Antibodies specific for this species, potentially recognizing the receptor-bound Aß tip ends and Conformation 1, may provide the opportunity to limit anti-amyloid ARIA toxicities associated with Aß plaque clearance.

## Methods

### Human brain tissue samples

Use of these pre-existing de-identified samples here was deemed exempt from ethics regulation by the Yale University Human Investigation Committee.The autopsy brain specimens were collected and fresh frozen at −80C at Yale University in New Haven, CT. By neuropathological criteria, the AD samples were each Braak stage V-VI, and CERAD C2-C3. The control brain was Braak stage 0I, and CERAD none.

### IC16 antibody design, expression, purification, and conjugation to HaloLink resin

The protein sequence of IC16 ScFv was previously published ^38^. A gene fragment based on reverse-translated IC16 ScFv amino acid sequence was synthesized by IDT technology, amplified by PCR and cloned into pcDNA3 mammalian expression vector as an N-terminal fusion to the HALO tag with IgG kappa signal peptide to achieve secreted expression. IC16 antibody was expressed using Expi293 Expression system and purified by nickel affinity chromatography. In brief, the media from transfected cells was bound in batch on nickel NTA resin (Qiagen). Resin was collected and washed with 20 bed volumes of phosphate buffered saline (PBS) then 20 bed volumes of PBS plus 10 mM imidazole. IC16 was eluted using PBS plus 250 mM Imidazole. Eluted IC16 antibody was pooled, concentrated to 1 mg/ml and dialyzed three times against PBS. The resulting purified antibody was stored at 4C, since we noted that freeze-thaw cycles have greatly reduced affinity.

To conjugate purified IC16 to a resin, the antibody was diluted to 0.25 mg/mL with PBS and mixed with HaloLink Resin (Promega) (1 mg of IC16 to 1 mL of resin) for at least four hours at room temperature. Conjugation was monitored by reading the unbound absorbance at 280 nm until >95% of the IC16 was conjugated to the resin. The resin was then collected and washed with 5 bed volumes of PBS + 0.001% DDM, 5 bed volumes of water, 1 bed volume of 0.2% ammonium hydroxide, and then immediately with 10 bed volumes PBS. IC16-conjugated HaloLink resin was stored in PBS with 0.01% sodium azide.

To prepare denatured IC-16 conjugated HaloLink resin used for preclearing, resin was washed with 3 bed volumes of 6M guanidine hydrochloride and then incubated with 3 bed volumes of 6M guanidine hydrochloride for 30 minutes at room temperature. After denaturation, the resin was washed thoroughly with at least 20 column volumes of PBS. This completely inactivates the IC16-ScFv, but leaves it covalently attached to the resin beads, allowing to use this resin for nonspecific binder absorption.

### Recombinant PrP expression and purification

Recombinant PrP^C^ was produced as described ^31, 39^. In short, the human PrP plasmid in pRSET-A vector containing an 6His-tag and a thrombin cleavage site was transformed in BL21(DE3) competent cells (NEB). Overnight growth of a starter culture in non-inducing medium MDAG135 was followed by a large-volume growth at room temperature in LB5052 autoinduction medium containing 1:1000 trace metals (Teknova, T1001), 2 mM MgSO_4_, 100 µg/mL ampicillin, 1x salt M and 1x 5052 autoinduction sugar mix ^40^. Cells were harvested by a 30-minute centrifugation at 4000 x *g* and lysed in Buffer G (6 M Guanidine HCl, 10 mM reduced glutathione, 100 mM Na_2_HPO_4_, 10 mM Tris-HCl, pH 8). The lysate was centrifuged at 17,700 x *g* for 90 min, and supernatant fraction was loaded onto HisPur Ni-NTA resin (Qiagen) in a gravity flow column. Protein was allowed to refold with a stepwise gradient of Buffer B (100 mM Na_2_HPO_4_, 10 mM Tris-HCl, pH 8) in Buffer G and eluted with 600 mM imidazole in 10 mM Na_2_HPO_4_, pH 5.8.

Eluted protein was dialyzed in 3.5 kDa Slide-A-Lyzer cassettes (ThermoFisher), in 5 mM Na_2_HPO_4_, pH 5.8 for 2 hours, and then overnight in water. His-tag was removed by overnight incubation with thrombin at 1 unit/mg. Subsequent cation exchange chromatography was performed using Source 15S column in AKTApure FPLC, equilibrated in 10 mM Na_2_HPO_4_, pH 6.5, and eluted in a 0-100% 20 CV gradient of 1M NaCl, 10 mM Na_2_HPO_4_, pH 6.5. Fractions were collected based on A280 in a single peak on the chromatogram and followed the same dialysis procedures.

### Preparation of synthetic Aß oligomers

For comparison with brain-derived receptor-bound Aß, samples of synthetic Aß42 oligomer as ADDLs (F12) and globulomer were prepared as described previously ^15, 31^.

### Immunoassays

To prepare the material for the epitope profiling studies by immunoblot both the purified receptor-bound Aß and sarkosyl-insoluble Aß fractions were lyophilized in screw-cap microcentrifuge tubes, then resuspended in HFIP and incubated for 2 h at 70C to monomerize the Aß assemblies. The tubes were then opened and kept in the fume hood overnight at room temperature to evaporate the HFIP. The remaining protein films were resuspended in 1X Laemmli buffer supplemented with 8 M urea. The resulting samples were separated on 4-20% gradient Criterion Tris-Glycine gels (BioRad), transferred onto the nitrocellulose membranes, blocked with Rockland fluorescent Western Blot blocking buffer and immunoprobed with the appropriate detection antibodies overnight at 4C, washed in PBST, incubated the IRdye conjugated secondary antibodies (LiCor) for 1 h at RT and imaged on the near-infrared scanner (LiCor Odyssey). Both primary and secondary antibody were diluted in Rockland fluorescent Western Blot blocking buffer. The immunoblot data quantitation was performed using the LiCor ImageStudio software.

To measure the levels of Aß assemblies in complex lysates as well as various steps of purification, several time-resolved fluorescence immunoassays were used. The general protocol has been previously published ^15^. White 384-well MaxiSorp polystyrene microplates (Thermofisher, 460372) were coated overnight at 4C with 20 µl/well of coating solution (30 mM Na_2_CO_3_, 80 mM NaHCO_3_, pH 9.6) containing 250 nM PrP^C^ (for PLISA assay), 100 nM IC16-Halo ScFv (for IC16–8243 assay) or 2 µg/ml Lecanemab (for Lecanemab–8243 assay). Following the coating the plates were washed with 3 dispensing/aspiration cycles of 100 µl/well of PBST (PBS, 0.05% Tween 20) using ELx405 microplate washer (BioTek) and blocked with 100 µl/well of Protein-Free T20 PBS blocking buffer (ThermoFisher, 37573) for 4 h at room temperature and then washed with PBST as above. Alongside with the tested samples, three-fold serial dilutions of ADDL Aßo ^13, 15, 17^ in F12 in PBSTB (PBST + 0.5% BSA) were applied to the coated plates in triplicates at 20 µl/well. The samples were diluted 2-fold with PBSTB and applied in triplicates. Additionally, three 4X dilutions were made for each sample to ensure linear quantitation in the samples that have widely different Aß levels. The plates were incubated with samples overnight at 4C, washed 3X with PBST and then incubated with D54D2 anti-Aß rabbit monoclonal antibody (Epitope in N terminus of Aß peptide, Cell Signaling Technology, #8243) diluted 1:2000 in PBSTB overnight at 4C. After 3X PBST wash to remove the unbound primary antibody, 20 µl/well of 1:8000 dilution of Eu-N1 Goat anti-Rabbit IgG (Aß and human lysate plates, Perkin-Elmer AD0105) in DELFIA assay buffer (Perkin Elmer 4002-0010) was applied for 2 h. Finally, for all the plates a final PBST wash was performed, 20 µl of DELFIA Enhancement Solution (Perkin Elmer 4001-0010) was added to each well, and time-resolved Europium fluorescence was measured using Victor 3V plate reader (Perkin Elmer).

### Receptor-bound Aβ purification from AD brain tissue

Post-mortem human brain slices were collected fresh frozen on dry ice and stored at −80C as described ^10^. For large scale purifications, approximately 80 g (wet weight) of brain was used. To prepare purified Aß from human brain, frozen frontal lobe cerebral cortex was minced and Dounce-homogenized on ice in four volumes of homogenization buffer 1X Tris buffered saline (TBS) supplemented with 1X complete EDTA free protease inhibitors (Roche). After homogenization, the homogenate volume was adjusted with homogenization buffer to result in 10 volumes of homogenization buffer per gram of brain. The homogenate was centrifuged for 30 minutes at 100,000 x *g* at 4C. The resulting pellet, P1, was then homogenized with 10 volumes of homogenization buffer and centrifuged again for 30 minutes at 100,000 x g at 4C. The resulting pellet, P2, was homogenized again in 10 volumes of 0.25X phosphate buffered saline (0.25X PBS) and centrifuged again for 30 minutes at 100,000 x *g* at 4C to generate a P3 pellet. For exhaustive extraction, the P3 pellet was homogenized again in 0.25X PBS as above and centrifuged an additional 3-5 times. Purified Aß used for cryo-EM and biological assays, the P3 pellet was homogenized in five volumes 0.25X PBS and five volumes of 1/4X PBS with 200 µM 20 kDa Poly(4-Styrenesulfonic acid-co-maleic acid) sodium salt (PSCMA, Sigma 434566-250G) was added (100 µM PSCMA final concentration) and incubated on a nutator for one hour at 4 C. This extract was then centrifuged for 60 min at 100,000 x *g* at 4C to generate supernatant S4 which contains PSCMA-extracted Aß species and P4 pellet. At that point the S4 supernatant and pellets were frozen and stored at −80C until further steps.

To purify and enrich receptor-bound Aß further, S4 was subjected to immunopurification. First S4 was precleared with denatured IC16 conjugated HALO resin on nutator for 4 h at 4C (1 ml resin to 80 ml of extract) to adsorb the proteins nonspecifically binding to the resin. After collecting the inactivated resin by centrifugation, the precleared supernatants were collected and IC-16 conjugated HALO resin was then added (1 ml resin to 40 ml of precleared S4) and mixed overnight at 4C to bind Aß to the resin. After binding, the resin was collected in a column and washed five column volumes (CV) of PBS. The resin was then transferred to 10 ml spin columns and washed further with another 5 CV PBS. Resin was washed 2×2 CV milliQ water via centrifugation. The resin was slightly dry after the second centrifugation step. Aß was eluted via centrifugation 3 times with one CV elution buffer (0.2% ammonium hydroxide). After each elution, the pH was neutralized using 0.3 volumes of 1% formic acid to pH 7.4. The resin was then washed thoroughly with PBS and the unbound was then bound and eluted for a second passage. After elution n-Dodecyl-beta-Maltoside (DDM) was added to 0.001% to prevent sample sticking to spin concentrators and the sample was concentrated 50-fold using 0.5 mL Amicon 50 kDa spin concentrators (Millipore). The resulting concentrate was aliquoted and flash frozen in liquid nitrogen and stored at −80C.

To generate sarkosyl-insoluble fibrils, the pellet P4 was extracted by a previously published protocol ^10^. In brief, the P4 pellet was homogenized in 20 volumes extraction buffer (10 mM Tris pH 7.5, 800 mM NaCl, 1 mM EGTA, 10% sucrose, and 1X complete EDTA free protease inhibitors). Sarcosyl was then added to 2% to the homogenate and incubated for 60 minutes shaking at 37C. This was then spun for 10 minutes at 10,000 x *g* at 25C and the supernatant was then subsequently spun at 100,000 x *g*. The pellet was then resuspended in extraction buffer at 1 ml/gram of initial P4 pellet. This was diluted three-fold with 50 mM Tris, 150 mM NaCl, 10% sucrose, and 0.2% Sarcosyl. This was first spun for 3 minutes at 3,000 x *g* and then supernatant was spun at 100,000 x *g* for 30 minutes. The resulting pellets were then resuspended and washed with resuspension buffer (20 mM HEPES pH 7.4, 50 mM NaCl) and collected by centrifugation at 18,000 x *g* for 30 min. This washing procedure was done three times. After the third wash, the pellets were resuspended in 0.1 ml suspension buffer per 1 gram of initial P4 pellet, aliquoted and flash frozen and stored at −80C.

### Sequential extractions from mouse brain tissue

APPswe/PSen1ΔE9 mice on WT or *Prnp*-/- background were previously described ^30^. The mice were housed and handled in compliance with IACUC regulations. Animals were sacrificed at 9-10 months of age and the extracted brains were frozen in liquid nitrogen and stored at −80C until use. To perform the biochemical brain fractionation the tissue was first Dounce-homogenized in 10 volumes of TBS. The resulting homogenates were centrifuged at 100,000 x *g* for 30 min, the TBS supernatants were saved and frozen at −80C and the pellets were resuspended in 10 volumes of TBS containing 1% Triton X100, incubated 30 min at 4C and centrifuged at 100,000 x *g* for 30 min. The TBS-TritonX supernatant was collected and the pellets were resuspended in 1/4X PBS and centrifuged at 100,000 x *g* for 30 min. The pellets resulting from this 1/4X PBS wash were resuspended in 30 µM PSCMA dissolved in 1/4X PBS, incubated 1 h at 4C and then centrifuged at 100,000 x *g* for 1 h. The resulting supernatant with PSCMA-extracted material was collected and frozen at −80C and the pellets were then used for extraction of insoluble proteins using 8 M urea, 4% SDS dissolved in TBS. The supernatants from TBS, TBS-Triton and PSCMA extractions were then analyzed using plate-based Aß assays.

### Synapse loss from human iPSC-derived cortical neurons

Human neurons derived from induced pluripotent stem cells (iPSCs) with Neurogenin2 (Ngn2) stably integrated under doxycycline control were prepared and cultured as described ^41^. Poly-D-Lysine-coated 384 well plates (Corning # 354663) were seeded with 2,000 cells in 50 µl of Classic Neuronal Media (CNR: 50% DMEM/F12 (Gibco), 50% Neurobasal-A (Gibco), 0.5X Glutamax (Gibco), 1X MEM Non-Essential Amino Acids Solution (Gibco), 0.5X B27 supplement (Gibco), 0.5X N-2 supplement (Gibco), 10 ng/ml BDNF (PeproTech), 10 ng/ml NT-3 (PeproTech), 1 µg/ml mouse Laminin (Gibco)) containing 2 µg/ml doxycycline hydrochloride (Sigma). On DIV 7 and 14, 50 µl of CNR without doxycycline hydrochloride was added to each well. Every week after that, 30 µl of media was removed from each well and replaced with 30 µL of fresh media. Cells were grown for at least 60 days before treatment with receptor-bound Aßo.

Receptor-bound Aß and the equivalent control brain fraction were prepared by dialyzing three times against 1,000X volume of Neurobasal-A using 100 µl dialysis cartridges (Thermo) at 4C. On the day of the experiment, 75% of the media was removed and mixed 1:1 with fresh CNR media and then Antibiotic-Antimitotic (Gibco) was added to 1X and filtered. The rest of the media was carefully flicked out of the plate and then 90 µl of the filtered media was added back to each well and the plates were returned to the 37C incubator for 2 h. After two hours, 10 µl of dialyzed Aß (with or without dilution), control brain, or dialysis media was added to the 384-well plate and the plate was returned to the incubator for 7 days.

After 7 days, plates were removed from the incubator, media was flicked out and replaced with 17 µl of fixation buffer (4% paraformaldehyde, 6% sucrose, and 1X PBS, and then returned to a 37C incubator for 30 min. The plates were washed two times with 1X PBS using a plate washer (Agilent) and 17 µl of blocking buffer (10 % horse serum (Gibco) in 1X PBST (PBS with 0.1% TritonX-100 (Sigma)) and incubated at room temperature for 30-60 minutes. Blocking buffer was removed and replaced with 17 µl of primary antibody solution (1% horse serum in PBST) containing 1:500 guinea pig anti-Synapsin 1/2 (Synaptic Systems # 106004), 1:500 rabbit anti-Homer1b/c (Synaptic Systems #160022), and 1:3000 chicken anti-MAP2 (Novus Biologics # NB300-213) and incubated overnight at 4C. Plates were then washed two times with 1X PBS and replaced with secondary antibody solution (1% horse serum in PBST) containing 1:500 goat anti-guinea pig Alexa Fluor 488 (Thermo), 1:500 goat anti-chicken Alexa Fluor 647 (Thermo), and 1:500 DAPI (Thermo) for 60 min at room temperature. Plates were then washed five times with 1X PBS and imaged. Cultures were imaged using ImageXpress Micro Confocal High-Content Imaging System (Molecular Devices) equipped with a 40X water objective.

Images were collected as maximum intensity projection of a 3.6 µm Z-stack with a 0.4 µm spacing. Image analysis was performed using ImageXpress built in analysis software, MetaXpress. First the MAP2 image was thresholded using an adaptive thresholding set to 1-10 µm and then the mask was expanded by 5 pixels. Then, both the Synapsin1/2 and Homer1b/c channels were run through a TopHat filter to even out the background signal and then through a find-round-objects filter to pick out the pre- and post-synaptic features for Synapsin1/2 and Homer1b/c, respectively. To measure synapsin1/2 density on MAP2, the number of synaptic puncta overlapping with the MAP2 mask were counted and then divided by the area of the MAP2 mask. To measure Synapsin1/2 colocalized with homer1b/c density on MAP2, first, only the round objects of Synapsin1/2 and Homer1b/c colocalized with each other that also overlap with MAP2 were counted and divided by the MAP2 area. To filter out images with either too few or too number of cells, total MAP2 for each image was averaged and images that were greater or less than two standard deviations from the mean were thrown out. Then the values for MAP2 area, Synapsin1/2 density on MAP2, and Synapsin1/2-Homer1b/c colocalized density on MAP2 for each image from the same well were averaged. Then fold change relative to the controls for each measurement was calculated as (Synaptic measurement (experimental condition))/(Synaptic measurement (control condition)).

### Immunogold negative staining

Six µl of brain-extracted receptor-bound Aß were mixed with 2 µl of recombinant PrP^C^ (50 nM) directly on a glow-discharged 300 mesh carbon-coated copper grid (Electron Microscopy Sciences) and allowed to incubate for 5 min at room temperature. After that, excess liquid from the grid surface was blotted off and the grid was floated onto a 40 µl paraformaldehyde (PFA) 2% for 2 minutes. Next, the carbon-coated side part of the grid was immediately floated on 40 µl droplet of blocking buffer deposited onto a Parafilm in a humidified chamber for 10 minutes. The blocking buffer consisted of a mixture of 0.2 g of BSA dissolved in 25 ml PBS with 25 µl of cold-water skin fish gelatin and 0.5 ml of 1% Tween20 (blocking buffer for TEM, Cold Spring Harbor Protocol). Subsequently, the excess solution was gently removed from the grid surface and the carbon-coated side of the grid was floated onto a 40 µl droplet of primary antibody, mouse anti-PrP (3F4, diluted 1:50 in blocking buffer) for 1 hour at room temperature in the same humidified chamber. Following that, the grid was washed with 5 drops of the blocking buffer (each washing was 2 minutes in 40 µl of blocking buffer), with the excess liquid blotted off between each wash. Subsequently, the grid was incubated with 6 nm gold-conjugated anti-mouse secondary antibody (diluted 1:20 in the blocking buffer) for 30 minutes at room temperature inside the humidified chamber. After five washes with Milli-Q water (each washing was 2 minutes) the grid was stained with 2 drops of 2% (w/v) uranyl acetate. The grid was briefly stained in the first drop, and in the second drop the grid was stained for 2 minutes. The sample was air-dried, and electron microscopy images were acquired on an FEI Tecnai T12 transmission electron microscope. The negative staining images revealed the presence of rPrP^C^ protein in close proximity to the tips of receptor-bound Aß. Filament length measurements for the location of gold-labeled PrP^C^ were done in Bsoft ^42^.

### Thioflavin-T polymerization assays

Fibril polymerization was monitored using Thioflavin-T (ThT) fluorescence similar to previous studies ^43, 44, 45, 46, 47, 48^. The polymerization reactions were prepared in a 96-well half-area clear-bottom plates (Corning) containing 100 µl/well. The plates were sealed with a plate sealer to prevent evaporation and incubated at 37C in the Victor Nivo multimode plate reader with continuous shaking (double orbital, 600 rpm). ThT fluorescence readings were recorded every 2 minutes for an interval of 36-48 h with excitation and emission wavelengths of 435 nm and 480 nm, respectively.

Prior to the polymerization reaction, synthetic Aß42 (The ERI Amyloid Laboratory, LLC) was monomerized by dissolving in 1,1,1,3,3,3-hexafluoro-2-propanol (HFIP) to 10 mg/ml, heated for two hours at 70C in sealed tubes, aliquoted into microfuge tubes (0.5 mg/tube), dried overnight, and then put under vacuum for one hour to remove residual HFIP. The HFIP-treated Aß42 film was first dissolved in 20 µl of anhydrous dimethyl sulfoxide (DMSO) and vortexed for 2 min to achieve a 5 mM Aß42 solution. This was diluted to 20 µM with 20 mM Hepes, 50 mM NaCl, pH 7.4 and immediately centrifuged at 14,000 x *g* for 5 min at 4C to remove insoluble aggregates. The resulting supernatant containing Aß42 monomer was used for polymerization reactions.

For the self-polymerization of synthetic Aß42 monomer, each 100 µl reaction mixture consisted of 5 µM synthetic Aß42 monomer and 10 µM ThT in polymerization buffer (20 mM Hepes, 50 mM NaCl, pH 7.4).

For seeding reactions with synthetic Aß42 fibril, the seeds were added in each well to generate a final concentration of 20 nM monomer equivalent concentration of Aß42 seeds equaling 0.4% of the unpolymerized Aß42 in the reaction. These seeds were derived from Aß filaments quiescently grown over 3 weeks at 23C from 100 µM Aß42 in 20 mM Hepes, 50 mM NaCl, pH 7.4. The resulting Aß42 filaments were diluted to 2 μM monomer equivalent concentration in a final volume of 2 ml and seeds were prepared by sonication using a probe sonicator (Branson) for 2 minutes on and 2 minutes off for a minimum of six times at 50% duty cycle and 25% output power. In control reactions, this sonication yielded filaments of 50-100 nm average length by negative stain EM.

For the seeding with sarkosyl-insoluble human Aß fibril seeds, the reaction mixture consisted of 5 µM final concentration of synthetic Aß42 monomer, 10 μM final concentration of ThT and appropriate dilutions of sarkosyl-insoluble human Aβ fibril seeds in each well. Prior to adding to the seeding reaction, the sarkosyl-insoluble human Aβ fibrils from Brain A and B were dialyzed against the polymerization buffer for at least 24 h. After dialysis, the sarkosyl-insoluble human Aβ fibrils were diluted to approximately 4 µM monomer equivalent concentration with polymerization buffer and seeds were generated using a probe sonicator (Branson) for 2 minutes on and 2 minutes off for a minimum of 6 times at 50% duty cycle and 25% output power. In control reactions, this sonication yielded filaments of 50-100 nm average length by negative stain EM.

For the seeding with Alzheimer’s purified receptor-bound Aß, the reaction mixture consisted of 5 µM final concentration of synthetic Aß42 monomer, 10 µM final concentration of ThT and 10-40 nM (monomer equivalent) final concentrations of receptor-bound Aß (from Brain A, B, C and D) and polymerization buffer in each 100 µl volume well. Prior to adding to the seeding reaction, the human receptor-bound Aß was dialyzed against the polymerization buffer for at least 24 h. The following control conditions for receptor-bound Aß effects were similarly dialyzed and found to have no effect on Aß42 monomer polymerization: (a) parallel preparations from control brain, (b) the vehicle buffer used for receptor-bound Aß, (c) 1.7 nM IC16 antibody, and (d) 5 nM PSCMA.

For the self-polymerization of synthetic Aß42 monomer in the presence of PrP^C^, the seeding reactions contained 5 µM synthetic Aß42 monomer, 400 nM recombinant PrP^C^ and 10 µM ThT in polymerization buffer. For seeding reactions with synthetic Aß42 fibril seeds and PrP^C^, the final concentrations were 5 µM Aß42 monomer, 30 nM monomer equivalent concentration of Aß42 seeds, 400 nM PrP^C^ and 10 μM ThT in polymerization buffer.

### Cryo-EM grid preparation and data acquisition

Four µl of the purified receptor-bound Aß or sarkosyl-insoluble Aß fibrils were applied on the carbon side of glow discharged (Edwards S150B) holey-carbon-coated grids (Quantifoil 300 mesh gold grids R 1.2/1.3, from Electron Microscopy Sciences). Subsequently, the sample was blotted using filter paper (Ø55/20 mm, Grade 595, Ted Pella) for 3-4 sec at a ‘blot force’ of 0 or ‘-1’ and plunge frozen in liquid ethane with a Vitrobot mark IV (Thermo). The sample chamber of the Vitrobot was maintained at 4C and a humidity level of 100%. The vitrified grids were screened on a Glacios transmission electron microscope (Thermo) operated at 200 keV and equipped with a K2 Summit direct electron detector. Electron micrograph movies for three-dimensional image reconstruction were acquired in super-resolution counting mode using a 300 keV Titan Krios electron microscope (Thermo) equipped with high brightness field emission gun (XFEG), a K3 direct electron detector (Gatan), and a GIF Quantum LS imaging energy filter (Gatan) (slit width of 20 eV) at the Yale Cryo-EM resources. The Serial-EM software (Hagen et al., 2017) was used for the automated collection of the micrograph movies at a nominal magnification of 81, (corresponding to a pixel size of 1.068 Å at the specimen level) with defocus range of −1.25 to −2.5 µm. All parameters corresponding to the cryo-EM data acquisition are presented in the Table 1.

### Cryo-EM data processing

Data sets were processed using standard amyloid helical reconstruction procedures implemented in RELION 4.0 ^49^. Dose-fractionated micrograph movies were corrected for beam-induced motion using the RELION’s own implementation of motion correction ^50^. Contrast transfer function (CTF) parameters were estimated from the motion-corrected micrographs using CTFFIND-4.1 ^51^. The start-end coordinates of the Aß filaments were manually marked in the aligned micrographs using RELION 4.0. Segments were extracted with an inter-box distance of 12-Å, a helical rise of 4.75 Å and using a box size of 200 pixels with 1.068 Å per pixel. Extracted segments were subjected to several rounds of reference free two-dimensional (2D) classification to remove suboptimal segments and to select for segments showing detailed features along the filament axis. A three-dimensional (3D) cylinder was generated using the “relion_helix_toolbox” utility and used as reference for an initial 3D classification of the helical segments retained after the 2D classification steps. The initial 3D classification performed on the set of optimal 2D-segments assumed a helical rise of 4.75 Å and a value of the helical twist as estimated from the crossover distances in the 2D class averages. Several rounds of 3D classification with optimization of helical twist and rise were performed to further select the best 3D classes for high-resolution 3D auto-refinement. Through 3D classification two conformations were separated for the Aß42 peptide monomer in the receptor-bound Aß and one conformation for Aß42 peptide monomer in the sarkosyl-insoluble Aß fibrils. A final round of 3D auto-refinement was performed after Bayesian polishing and per-particle CTF refinement ^50, 52^. The final reconstructions were post-processed with a soft-edge mask and B-factor sharpened. The resolution of each 3D reconstructions was estimated from the value of the Fourier Shell Correlation (FSC) curve at 0.143 cutoff between two independently refined half-maps using phase randomization to correct for convolution effects of a generous, soft-edged solvent mask ^53^. Details about cryo-EM data processing are presented in Table 1.

### Model building and refinement, structural analysis and visualization

Initial models were generated by rigid body fitting of PDB entry 7Q4B ^10^ into the postprocessed electron microscopy maps of receptor-bound Aß (Conformation 1, Brain A), receptor-bound Aß (Conformation 2, Brain B) and sarkosyl-insoluble Aß fibrils (Brain A). Rigid body fitting was performed using the fit-in-map tool in Chimera ^54^. Subsequently, the atomic models were manually built and adjusted in Coot iteratively ^55^ and refined using real-space-refinement implemented in PHENIX ^56^ for five macro-cycles with applied secondary structure, rotamer, and Ramachandran restraints. The models were validated using MolProbity ^57^ and cryo-EM validation tools within PHENIX. Figures were prepared using UCSF Chimera and ChimeraX ^58^.

## Data availability

The cryo-EM maps generated in this study have been deposited in the Electron Microscopy Data Bank under the accession codes EMD-45642 (sarkosyl-insoluble Aß42 filaments), EMD-45647 (receptor-bound Aß42, conformation 2) and EMD-45770 (receptor-bound Aß42, conformation 1). The structural model generated in this study have been deposited in the Protein Data Bank under the accession codes PDB ID 9CK6 (sarkosyl-insoluble Aß42 filaments), PDB ID 9CKI (receptor-bound Aß42, Conformation 2) and PDB ID 9CO4 (receptor-bound Aß42, Conformation 1).

## Acknowledgments

We thank Jian Liu for excellent technical assistance, and Dr. Shenping Wu for Titan Krios data collection at the Yale West Campus Cryo-EM facility. We also thank Dr. Marc Llaguno, and Dr. Xinran Liu for managing the microscope resources at the Center for Cellular and Molecular Imaging, Electron Microscopy Facility at Yale Medical School. This work was supported by National Institutes of Health grants R01AG034924, RF1AG070926 and P30AG066508 to SMS and T32NS105583 to GPR.

## Author Contributions

Conceptualization: MAK, CB, GPR, SMS; Methodology: MAK, CB, GPR, YL, PG; Investigation: MAK, CB, GPR, YL, PG; Visualization: MAK, CB, GPR; Funding acquisition: SMS; Project administration: SMS; Supervision: SMS; Writing – original draft: MAK, CB, GPR, SMS; Writing – review & editing: MAK, CB, GPR, YL, PG, SMS

## Extended data

**Extended Data Fig. 1.**
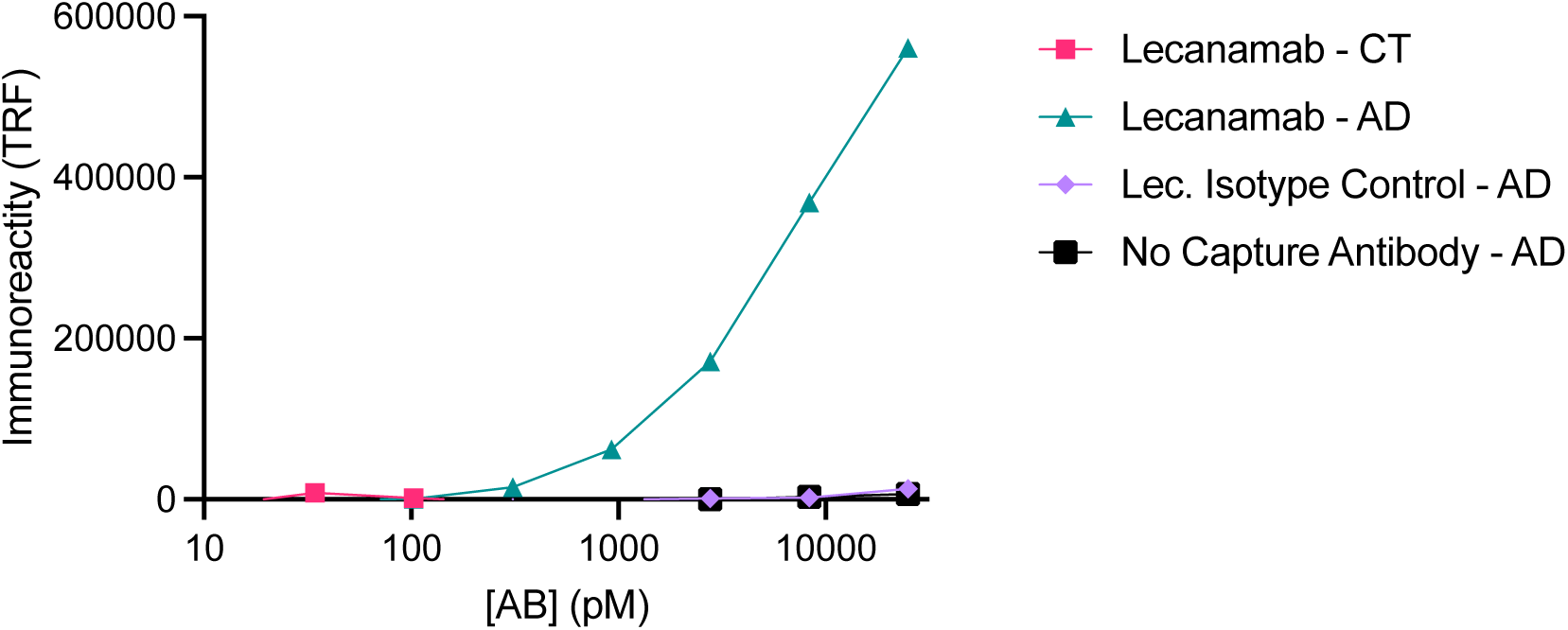
Brain purified receptor-bound Aß reacts with Lecanemab. Brain purified receptor-bound Aß binds to ELISA plates coated with 5 µg/mL Lecanamab but not to wells coated with isotype control or to no antibody wells. The equivalent fraction from control brain (CT) has undetectable levels of Aß immunoreactivity.

**Extended Data Fig. 2.**
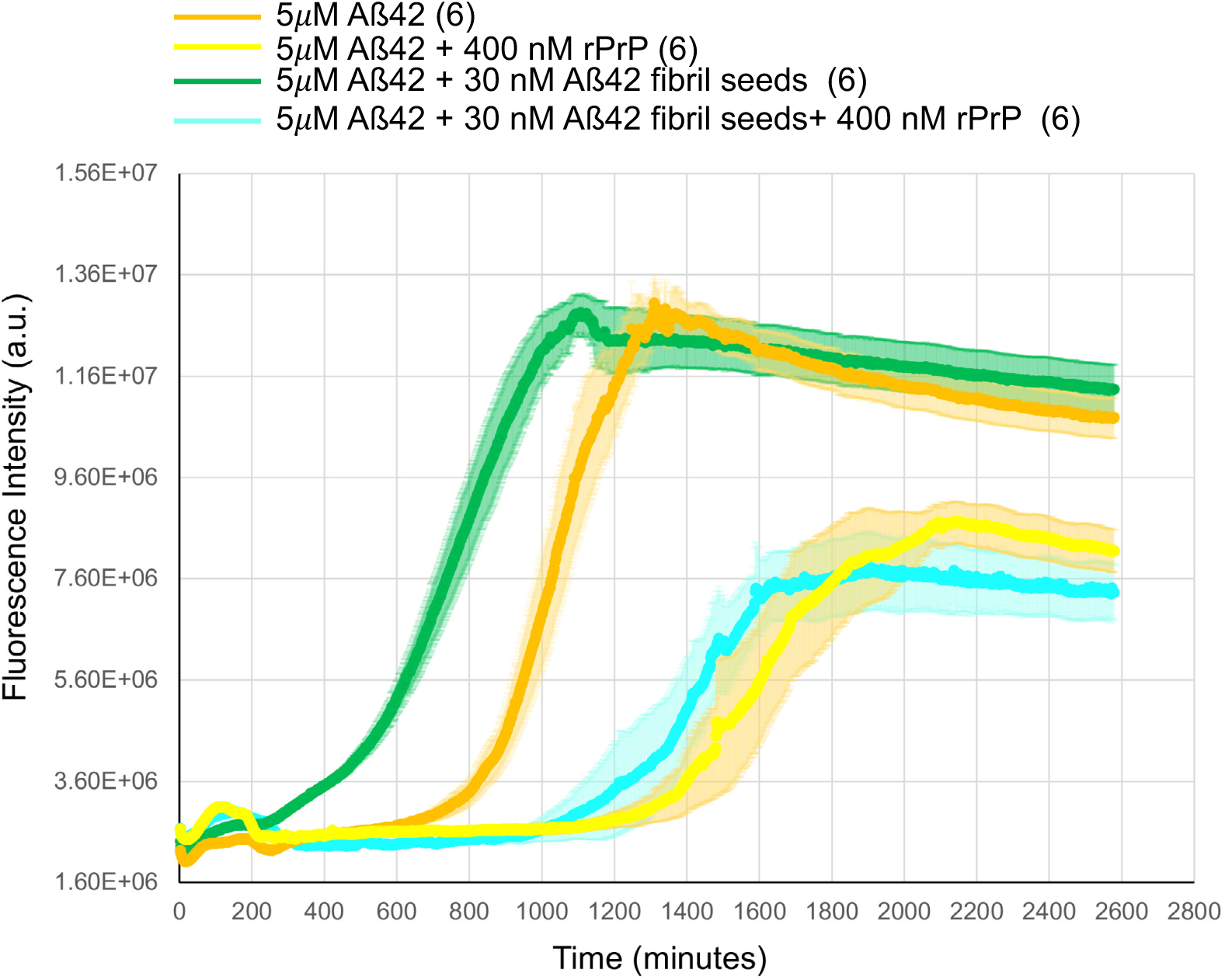
Recombinant excess PrP^C^ inhibits Aß42 amyloid filament formation. Thioflavin T fluorescence measurements as a function of time for 5 µM synthetic Aß42 polymerized alone (orange), or with 30 nM (monomer equivalent) synthetic Aß42 fibril seeds (green), or with 400 nM PrP^C^ (yellow), or with 30 nM synthetic Aß42 fibril seeds plus 400 nM PrP^C^ (cyan). Error bars represent means ± SEM for N=6.

**Extended Data Fig. 3.**
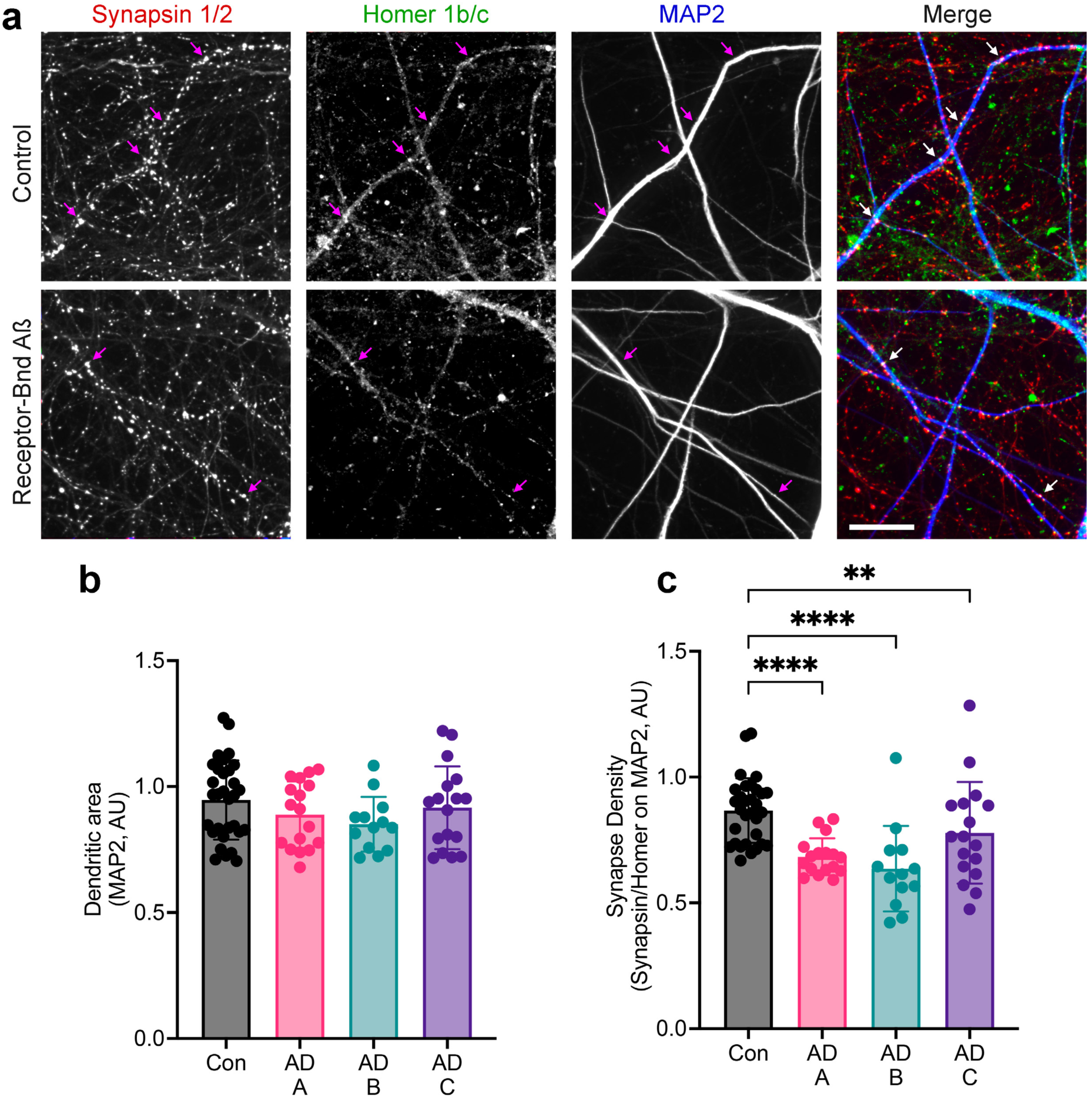
Purified receptor-bound Aß potently suppresses synapse density in human iPSC derived neurons. **a,** Human iPSC cultures were differentiated to cortical excitatory neurons by dox-induced expression of Neurogenin2. After 60-65 days of culture, the neurons were exposed to purified receptor-bound Aß (Brain A-C) or equivalent fraction from control brain (CT) for seven days prior to fixation and staining. The neurons were exposed to a final concentration of AD brain purified receptor-bound Aß of 160 pM Aß for Brain A, 80 pM for Brain B and 16 pM for Brain C (40 nM Aß monomer equivalent for Brain A, 20 nM for B and 4 nM for C). Equivalent fractions from control brain contained no detectable Aß. Fixed neurons were stained with antibodies reactive to MAP2, Synapsin1/2, and Homer1b/c. Images of individual channels and merged image collected with an automated 384 well spinning disc confocal microscope system channels are shown. The purple arrows indicated colocalization of Synapsin and Homer within MAP2 positive area detected by image analysis. Scale bar, 36 µm. **b,** Dendrite density detected as MAP2 area from images as in **a**. Dendrite does not change after addition of purified receptor-bound Aß versus equivalent fraction from control brain. Error bars represent the mean ± SEM. Each dot is the average value of 2-4 images from separate cultures. No statistically significant differences detected. **c,** Synaptic puncta identified as Synapsin/Homer double positive regions within MAP2 area as in **a**. Mean ± SEM, each dot is the average value of 2-4 images from separate cultures. ****, P<0.0001; **, P<0.01 one way ANOVA mixed effect model with matching and Dunnett correction for multiple tests.

**Extended Data Fig. 4.**
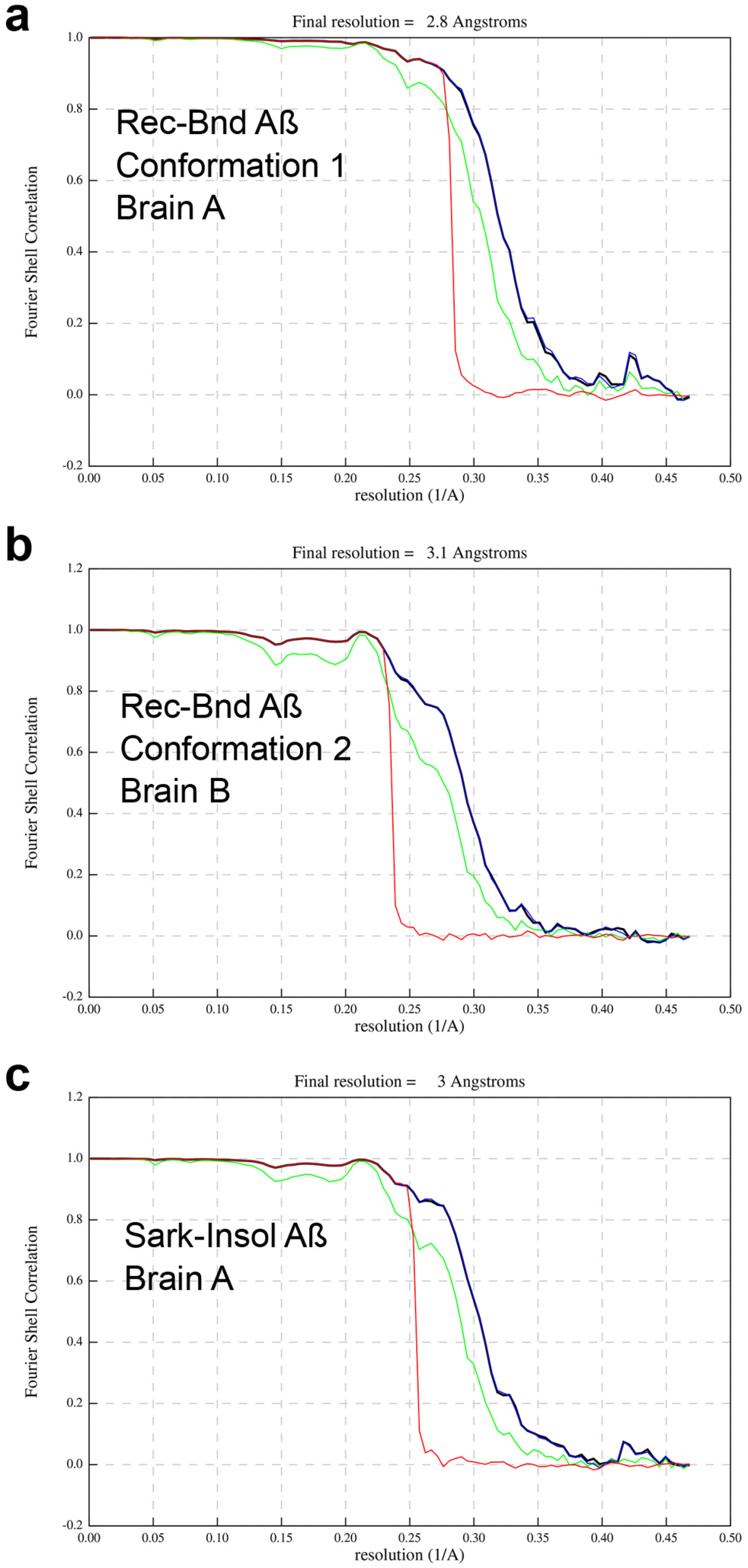
Fourier Shell Correlation (FSC) traces for each reconstruction. **a,** FSC traces between the two half-maps for the receptor-bound Aß in Conformation 1, from Brain A. **b,** FSC traces between the two half-maps for the receptor-bound Aß in Conformation 2, from Brain B. **c,** FSC traces between the two half-maps for the sarkosyl-insoluble human Aß42 filaments from Brain A. Black trace, FSC corrected. Green trace, FSC unmasked maps. Blue, FSC masked maps. Red trace, corrected FSC phase randomized masked maps.

**Extended Data Fig. 5.**
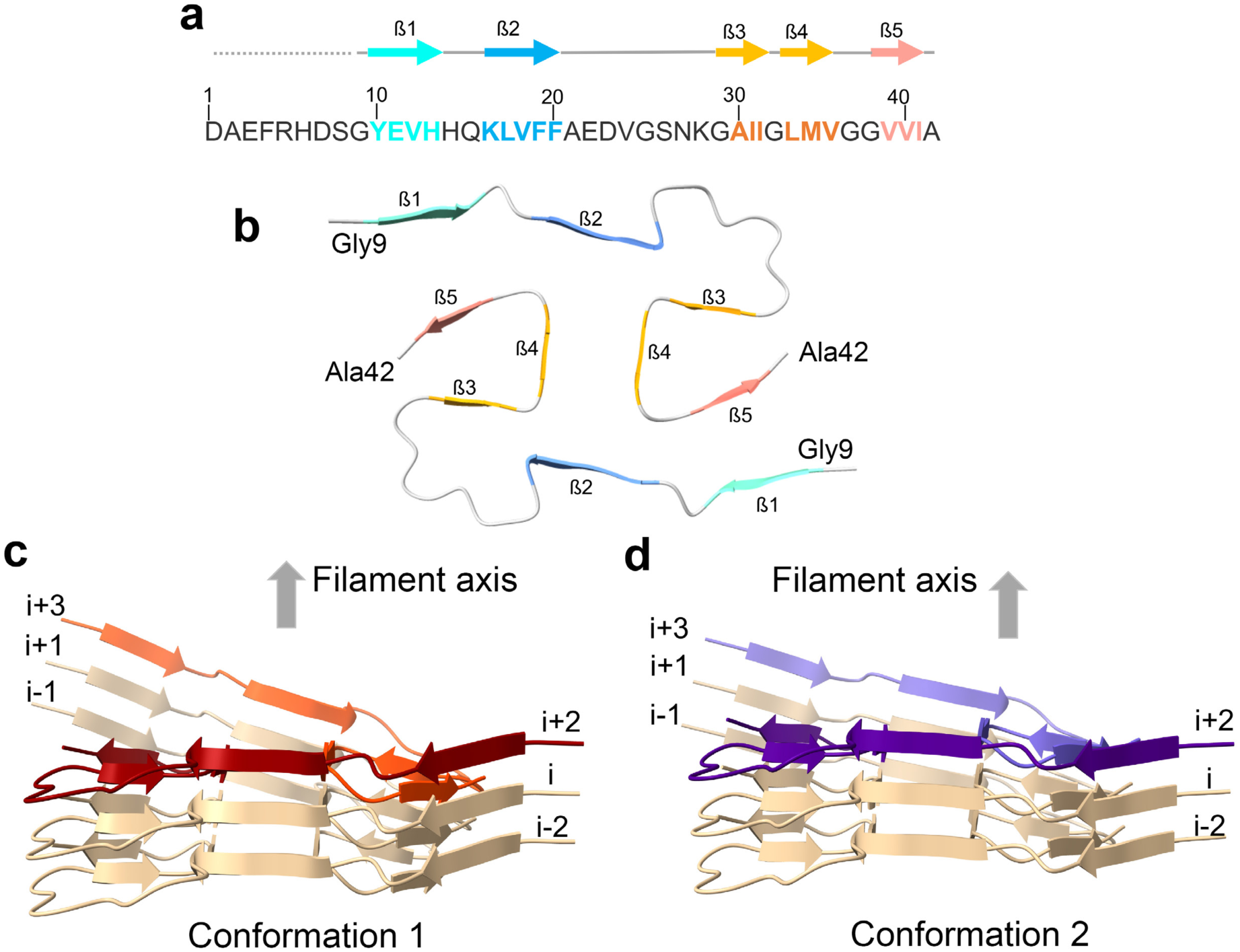
Juxtaposition of Aß monomers from different rungs of receptor-bound Aß Conformations. **a,** Primary sequence of Aß42 with secondary structure assignment of ß sheet conformation to specific residues for receptor-bound Aß Conformations 1 and 2. **b,** Ribbon diagram showing two monomers of Aß9-42 within one rung viewed from the long axis of the receptor-bound Aß rod. **c, d,** Ribbon diagrams of 6 Aß peptide structures from 3 rungs in Conformation 1 (**c**) versus Conformation 2 (**d**). Note that in Conformation 1 where the amino terminus of one peptide (i+2) is juxtaposed to the carboxyl terminus of the opposite peptide (i+3), the carboxyl terminus of that same peptide (i+2) is adjacent to amino terminus of the opposite peptide two rungs below (i-1). In contrast, the displacement in Conformation 2 is one rung (i+1), rather than two.

**Extended Data Fig. 6.**
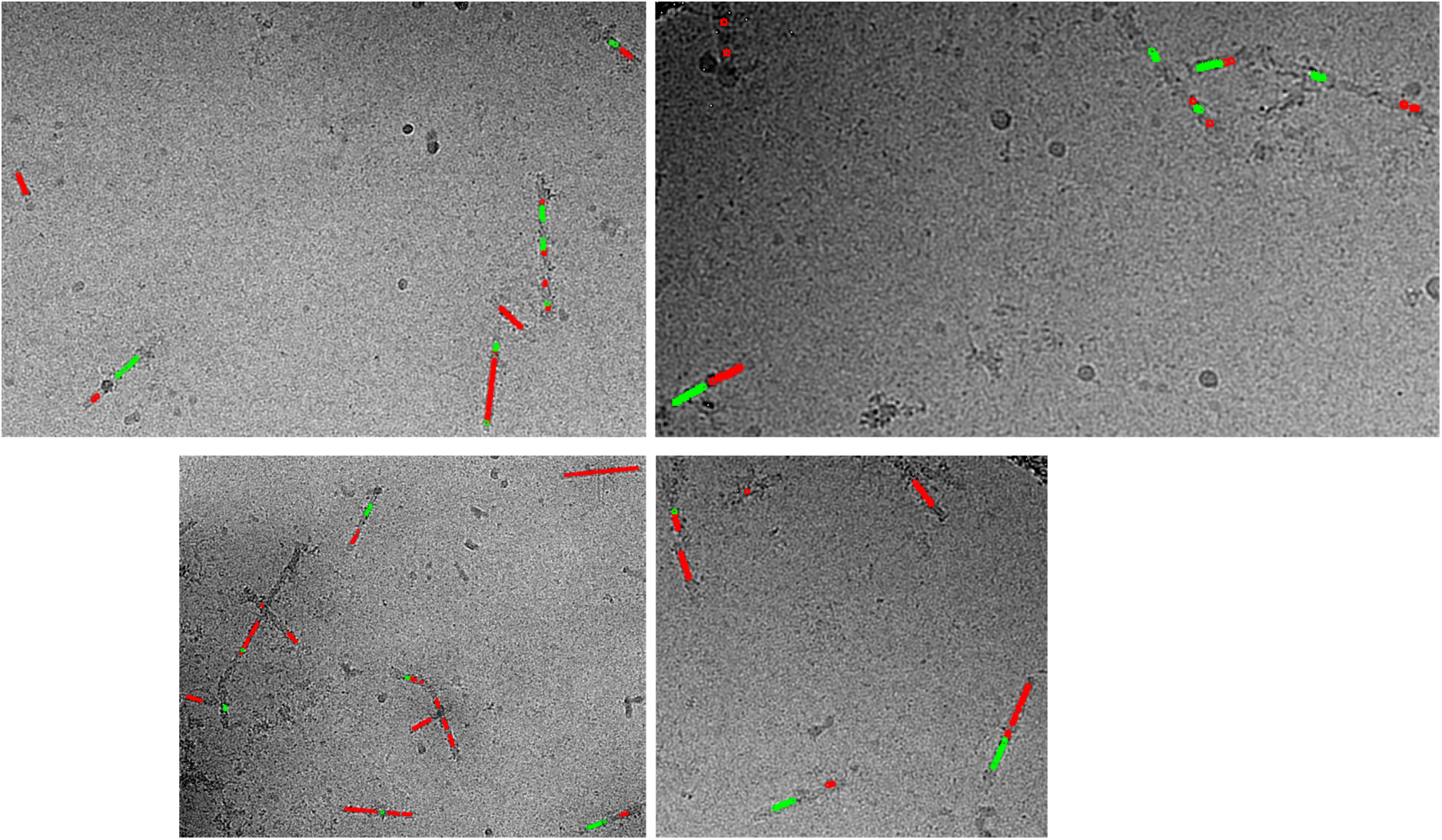
Distribution of Aß conformations along individual receptor-bound Aß rods. Structural variation across the length of the PSCMA-extractable, receptor-bound Aß isolated from Brain A. Tracing of the two conformations on individual Aß filaments (from Brain A) in cryo-EM micrographs. Red, Conformation 1; green, Conformation 2.

**Extended Data Fig. 7.**
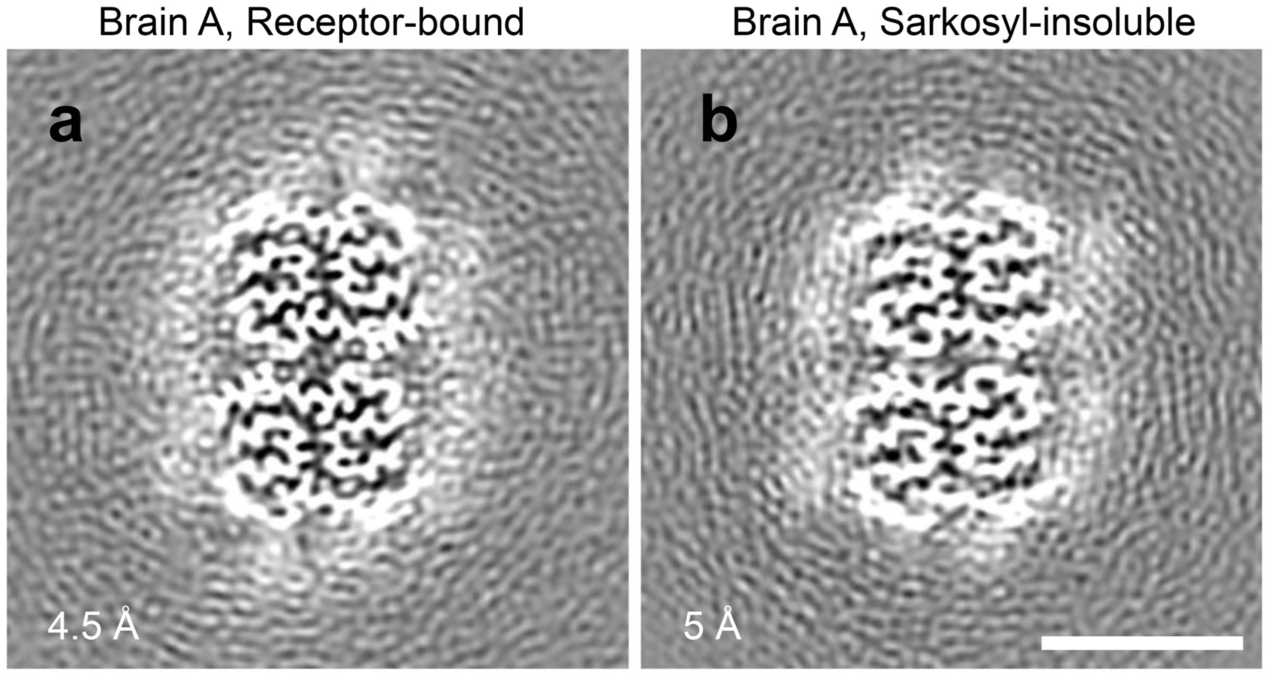
Rare doublets composed of Conformation 2 Aß peptide. **a,** Aß filament doublet from the Brain A receptor-bound Aß sample; resolved at 4.5 Å. **b,** Aß filament doublet from the Brain A sarkosyl-insoluble sample; resolved at 5.0 Å. Bar, 5 nm.

**Extended Data Fig. 8.**
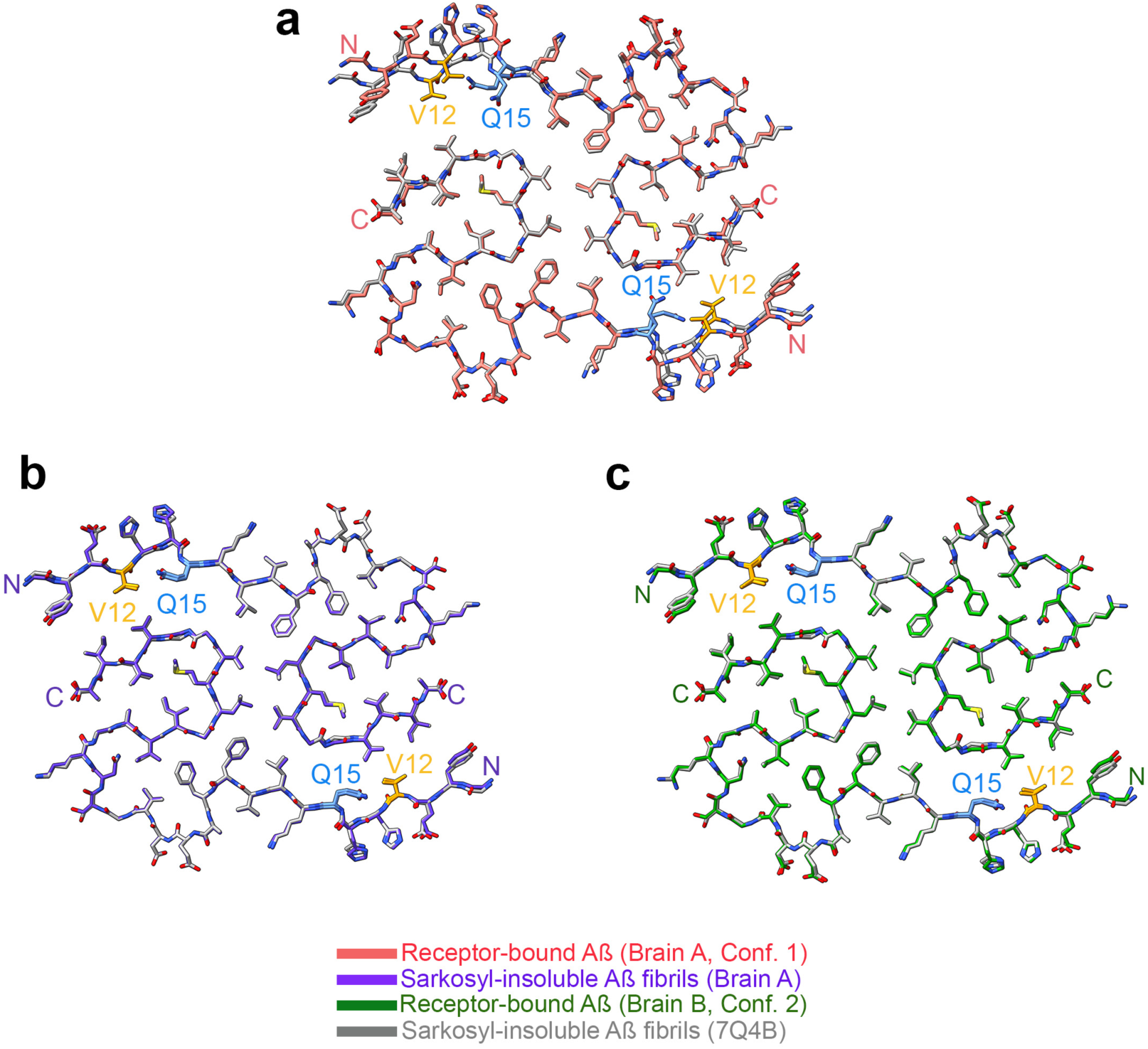
Comparison of receptor-bound Aß with published Aß fibril conformation. **a,** Atomic model of one rung of the receptor-bound Aß from Brain A (Conformation 1) superposed with the atomic model of one rung of the sarkosyl-insoluble Aß filaments (pdb:7Q4B) with an r.m.s.d. of 1.52 Å over 68 C𝛼 atoms. **b,** Atomic model of one rung of the Aß filaments from sarkosyl–insoluble Brain A superposed with the atomic model of one rung of the sarkosyl-insoluble Aß filaments (pdb:7Q4B) with an r.m.s.d. of 0.265 Å over 68 C𝛼 atoms. **c,** Atomic model of one rung of the receptor-bound Aß from Brain B (Conformation 2) superposed with the atomic model of one rung of the sarkosyl-insoluble Aß filaments (pdb:7Q4B) with an r.m.s.d. of 0.22 Å over 68 C𝛼 atoms.

**Extended Data Fig. 9.**
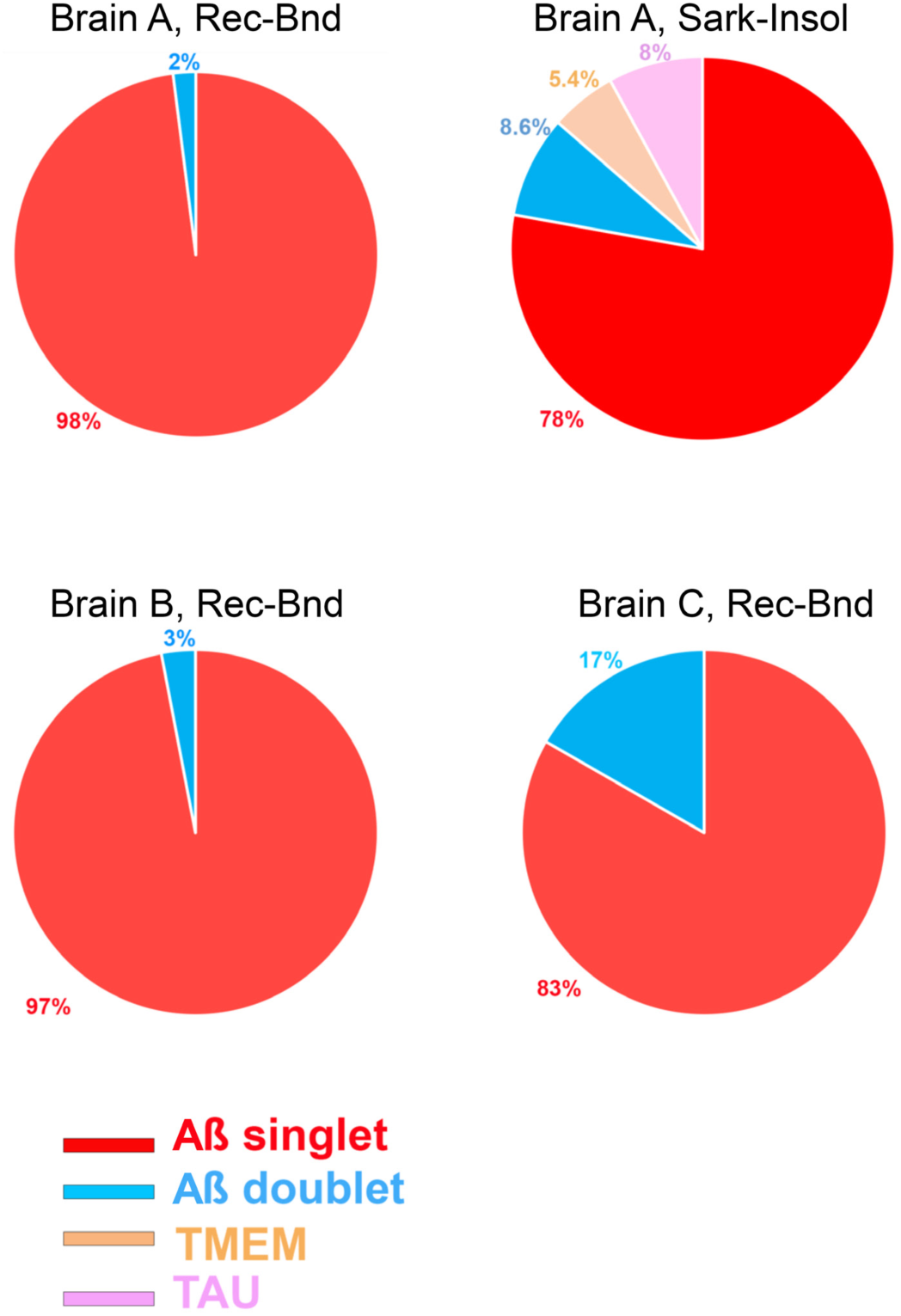
Sarkosyl-insoluble extracts of AD brain contain Aß, Tau and TMEM106B fibrils. Proportion of different amyloid assemblies observed in the various brain preparations.

## References

1. Knopman DS, et al. Alzheimer disease. Nat Rev Dis Primers 7, 33 (2021).

2. Long JM, Holtzman DM. Alzheimer Disease: An Update on Pathobiology and Treatment Strategies. Cell 179, 312–339 (2019).

3. van Dyck CH, et al. Lecanemab in Early Alzheimer’s Disease. N Engl J Med 388, 9–21 (2023).

4. Sims JR, et al. Donanemab in Early Symptomatic Alzheimer Disease: The TRAILBLAZER-ALZ 2 Randomized Clinical Trial. JAMA : the journal of the American Medical Association 330, 512–527 (2023).

5. Petersen RC, et al. NIA-AA Alzheimer’s Disease Framework: Clinical Characterization of Stages. Ann Neurol 89, 1145–1156 (2021).

6. Selkoe DJ, Hardy J. The amyloid hypothesis of Alzheimer’s disease at 25 years. EMBO Mol Med 8, 595–608 (2016).

7. Hampel H, et al. The Amyloid-beta Pathway in Alzheimer’s Disease. Mol Psychiatry 26, 5481–5503 (2021).

8. Walsh DM, et al. Naturally secreted oligomers of amyloid beta protein potently inhibit hippocampal long-term potentiation in vivo. Nature 416, 535–539 (2002).

9. Benilova I, Karran E, De Strooper B. The toxic Abeta oligomer and Alzheimer’s disease: an emperor in need of clothes. Nat Neurosci 15, 349–357 (2012).

10. Yang Y, et al. Cryo-EM structures of amyloid-beta 42 filaments from human brains. Science 375, 167–172 (2022).

11. Zielinski M, et al. Cryo-EM of Abeta fibrils from mouse models find tg-APP(ArcSwe) fibrils resemble those found in patients with sporadic Alzheimer’s disease. Nat Neurosci 26, 2073–2080 (2023).

12. Yang Y, et al. Cryo-EM structures of amyloid-beta filaments with the Arctic mutation (E22G) from human and mouse brains. Acta Neuropathol 145, 325–333 (2023).

13. Lambert MP, et al. Diffusible, nonfibrillar ligands derived from Abeta1-42 are potent central nervous system neurotoxins. Proc Natl Acad Sci U S A 95, 6448–6453 (1998).

14. Yu L, et al. Structural characterization of a soluble amyloid beta-peptide oligomer. Biochemistry 48, 1870–1877 (2009).

15. Kostylev MA, et al. Prion-Protein-interacting Amyloid-beta Oligomers of High Molecular Weight Are Tightly Correlated with Memory Impairment in Multiple Alzheimer Mouse Models. J Biol Chem 290, 17415–17438 (2015).

16. Stern AM, et al. Abundant Abeta fibrils in ultracentrifugal supernatants of aqueous extracts from Alzheimer’s disease brains. Neuron 111, 2012–2020 e2014 (2023).

17. Lauren J, Gimbel DA, Nygaard HB, Gilbert JW, Strittmatter SM. Cellular prion protein mediates impairment of synaptic plasticity by amyloid-beta oligomers. Nature 457, 1128–1132 (2009).

18. Smith LM, Kostylev MA, Lee S, Strittmatter SM. Systematic and standardized comparison of reported amyloid-beta receptors for sufficiency, affinity, and Alzheimer’s disease relevance. J Biol Chem 294, 6042–6053 (2019).

19. Smith LM, Strittmatter SM. Binding Sites for Amyloid-beta Oligomers and Synaptic Toxicity. Cold Spring Harbor perspectives in medicine 7, (2017).

20. Purro SA, Nicoll AJ, Collinge J. Prion Protein as a Toxic Acceptor of Amyloid-beta Oligomers. Biol Psychiatry 83, 358–368 (2018).

21. Stoner A, et al. Neuronal transcriptome, tau and synapse loss in Alzheimer’s knock-in mice require prion protein. Alzheimer’s research & therapy 15, 201 (2023).

22. Spurrier J, et al. Reversal of synapse loss in Alzheimer mouse models by targeting mGluR5 to prevent synaptic tagging by C1Q. Science translational medicine 14, eabi8593 (2022).

23. Gunther EC, et al. Rescue of Transgenic Alzheimer’s Pathophysiology by Polymeric Cellular Prion Protein Antagonists. Cell Rep 26, 145–158 e148 (2019).

24. Cox TO, et al. Anti-PrP(C) antibody rescues cognition and synapses in transgenic alzheimer mice. Ann Clin Transl Neurol 6, 554–574 (2019).

25. Salazar SV, et al. Conditional Deletion of Prnp Rescues Behavioral and Synaptic Deficits after Disease Onset in Transgenic Alzheimer’s Disease. J Neurosci 37, 9207–9221 (2017).

26. Heiss JK, Barrett J, Yu Z, Haas LT, Kostylev MA, Strittmatter SM. Early Activation of Experience-Independent Dendritic Spine Turnover in a Mouse Model of Alzheimer’s Disease. Cereb Cortex 27, 3660–3674 (2017).

27. Haas LT, et al. Silent Allosteric Modulation of mGluR5 Maintains Glutamate Signaling while Rescuing Alzheimer’s Mouse Phenotypes. Cell Rep 20, 76–88 (2017).

28. Haas LT, Salazar SV, Kostylev MA, Um JW, Kaufman AC, Strittmatter SM. Metabotropic glutamate receptor 5 couples cellular prion protein to intracellular signalling in Alzheimer’s disease. Brain 139, 526–546 (2016).

29. Um JW, et al. Metabotropic glutamate receptor 5 is a coreceptor for Alzheimer abeta oligomer bound to cellular prion protein. Neuron 79, 887–902 (2013).

30. Gimbel DA, et al. Memory impairment in transgenic Alzheimer mice requires cellular prion protein. J Neurosci 30, 6367–6374 (2010).

31. Kostylev MA, et al. Liquid and Hydrogel Phases of PrP(C) Linked to Conformation Shifts and Triggered by Alzheimer’s Amyloid-beta Oligomers. Molecular cell 72, 426–443 e412 (2018).

32. Soderberg L, et al. Lecanemab, Aducanumab, and Gantenerumab - Binding Profiles to Different Forms of Amyloid-Beta Might Explain Efficacy and Side Effects in Clinical Trials for Alzheimer’s Disease. Neurotherapeutics : the journal of the American Society for Experimental NeuroTherapeutics 20, 195–206 (2023).

33. Amin L, Harris DA. Abeta receptors specifically recognize molecular features displayed by fibril ends and neurotoxic oligomers. Nat Commun 12, 3451 (2021).

34. Nieznanski K, Surewicz K, Chen S, Nieznanska H, Surewicz WK. Interaction between prion protein and Abeta amyloid fibrils revisited. ACS chemical neuroscience 5, 340–345 (2014).

35. Fernandez A, et al. Cryo-EM structures of amyloid-beta and tau filaments in Down syndrome. Nat Struct Mol Biol 31, 903–909 (2024).

36. Lesne S, et al. Retraction Note: A specific amyloid-beta protein assembly in the brain impairs memory. Nature 631, 240 (2024).

37. Demattos RB, et al. A plaque-specific antibody clears existing beta-amyloid plaques in Alzheimer’s disease mice. Neuron 76, 908–920 (2012).

38. Dornieden S, et al. Characterization of a single-chain variable fragment recognizing a linear epitope of abeta: a biotechnical tool for studies on Alzheimer’s disease? PLoS One 8, e59820 (2013).

39. Zahn R, von Schroetter C, Wuthrich K. Human prion proteins expressed in Escherichia coli and purified by high-affinity column refolding. FEBS Lett 417, 400–404 (1997).

40. Studier FW. Protein production by auto-induction in high density shaking cultures. Protein Expr Purif 41, 207–234 (2005).

41. Tian R, et al. CRISPR Interference-Based Platform for Multimodal Genetic Screens in Human iPSC-Derived Neurons. Neuron 104, 239–255 e212 (2019).

42. Heymann BJ. Bsoft: Image Processing for Structural Biology. Bio Protoc 12, e4393 (2022).

43. Liang R, Tian Y, Viles JH. Cross-seeding of WT amyloid-beta with Arctic but not Italian familial mutants accelerates fibril formation in Alzheimer’s disease. J Biol Chem 298, 102071 (2022).

44. Ruiz-Riquelme A, et al. Abeta43 aggregates exhibit enhanced prion-like seeding activity in mice. Acta neuropathologica communications 9, 83 (2021).

45. Koloteva-Levine N, et al. Amyloid particles facilitate surface-catalyzed cross-seeding by acting as promiscuous nanoparticles. Proc Natl Acad Sci U S A 118, (2021).

46. Korshavn KJ, et al. Reduced Lipid Bilayer Thickness Regulates the Aggregation and Cytotoxicity of Amyloid-beta. J Biol Chem 292, 4638–4650 (2017).

47. Salvadores N, Shahnawaz M, Scarpini E, Tagliavini F, Soto C. Detection of misfolded Abeta oligomers for sensitive biochemical diagnosis of Alzheimer’s disease. Cell Rep 7, 261–268 (2014).

48. Hellstrand E, Boland B, Walsh DM, Linse S. Amyloid beta-protein aggregation produces highly reproducible kinetic data and occurs by a two-phase process. ACS chemical neuroscience 1, 13–18 (2010).

49. Kimanius D, Dong L, Sharov G, Nakane T, Scheres SHW. New tools for automated cryo-EM single-particle analysis in RELION-4.0. Biochem J 478, 4169–4185 (2021).

50. Zivanov J, Nakane T, Scheres SHW. A Bayesian approach to beam-induced motion correction in cryo-EM single-particle analysis. IUCrJ 6, 5–17 (2019).

51. Rohou A, Grigorieff N. CTFFIND4: Fast and accurate defocus estimation from electron micrographs. J Struct Biol 192, 216–221 (2015).

52. Zivanov J, Nakane T, Scheres SHW. Estimation of high-order aberrations and anisotropic magnification from cryo-EM data sets in RELION-3.1. IUCrJ 7, 253–267 (2020).

53. Chen S, et al. High-resolution noise substitution to measure overfitting and validate resolution in 3D structure determination by single particle electron cryomicroscopy. Ultramicroscopy 135, 24–35 (2013).

54. Pettersen EF, et al. UCSF Chimera--a visualization system for exploratory research and analysis. J Comput Chem 25, 1605–1612 (2004).

55. Emsley P, Cowtan K. Coot: model-building tools for molecular graphics. Acta Crystallogr D Biol Crystallogr 60, 2126–2132 (2004).

56. Afonine PV, et al. Real-space refinement in PHENIX for cryo-EM and crystallography. Acta Crystallogr D Struct Biol 74, 531–544 (2018).

57. Prisant MG, Williams CJ, Chen VB, Richardson JS, Richardson DC. New tools in MolProbity validation: CaBLAM for CryoEM backbone, UnDowser to rethink “waters,” and NGL Viewer to recapture online 3D graphics. Protein Sci 29, 315–329 (2020).

58. Pettersen EF, et al. UCSF ChimeraX: Structure visualization for researchers, educators, and developers. Protein Sci 30, 70–82 (2021).

